# Rpb4 and Puf3 imprint and post-transcriptionally control the stability of a common set of mRNAs in yeast

**DOI:** 10.1101/2020.07.25.220095

**Authors:** A.I. Garrido-Godino, I. Gupta, F. Gutiérrez-Santiago, A.B. Martínez-Padilla, A. Alekseenko, L.M. Steinmetz, J.E. Pérez-Ortín, V. Pelechano, F. Navarro

## Abstract

Gene expression involving RNA polymerase II is regulated by the concerted interplay between mRNA synthesis and degradation, crosstalk in which mRNA decay machinery and transcription machinery respectively impact transcription and mRNA stability. Rpb4, and likely dimer Rpb4/7, seem the central components of the RNA pol II governing these processes. In this work we unravel the molecular mechanisms participated by Rpb4 that mediate the posttranscriptional events regulating mRNA imprinting and stability. By RIP-Seq, we analyzed genome-wide the association of Rpb4 with mRNAs and demonstrated that it targeted a large population of more than 1400 transcripts. A group of these mRNAs was also the target of the RNA binding protein, Puf3. We demonstrated that Rpb4 and Puf3 physically, genetically, and functionally interact and also affect mRNA stability, and likely the imprinting, of a common group of mRNAs. Furthermore, the Rpb4 and Puf3 association with mRNAs depends on one another. We also demonstrated, for the first time, that Puf3 associates with chromatin in an Rpb4-dependent manner. Our data also suggest that Rpb4 could be a key element of the RNA pol II that coordinates mRNA synthesis, imprinting and stability in cooperation with RBPs.

## INTRODUCTION

Gene expression is a highly regulated process that comprises several coordinated steps to ensure appropriate mRNA levels, and is determined by two different processes: mRNA synthesis and decay. Both processes occur mainly in different cell locations and need a high coordination. Changes in mRNA levels are determined by their transcription and degradation rates, and the parallel co-regulation of global mRNA synthesis and decay is important to maintain mRNA homeostasis ^1–4^.

mRNA imprinting, defined by M. Choder as the co-transcriptional association of a protein to an mRNA that lasts throughout the mRNA lifetime and is required for proper regulation of all major stages that the mRNA undergoes ^5^, is crucial for coordinating transcription in the nucleus and the decay that occurs mainly in the cytoplasm ^1, 5–11^. Two RNA polymerase II subunits, Rpb4 and Rpb7, correspond to these mRNA imprinting proteins interconnecting both processes.

Rpb4 and Rpb7 bind mRNA in the RNA pol II context during transcription and remain associated with mRNA throughout its life by regulating processes such as export, translation and decay ^9, 12-16^. Rpb4 has been proposed to interact with mRNA in an Rpb7 dependent manner ^7, 9, 17, 18^. The Rpb4/7 dimer participates in several transcription steps, such as transcription initiation ^19^ and elongation ^20^, and dissociates from the RNA pol II core by interacting with mRNA during transcription ^7, 10, 13^, although this statement is not unanimously accepted ^7, 9, 17, 18, 21^. Furthermore Rpb4 participates in transcription termination and gene looping ^22^. Rpb4-mediated mRNA decay probably occurs by interacting with mRNA decay machinery elements, such as Pat1-Lsm1/7 ^14^. Under optimal growth conditions, Rpb4 serves as a key protein to globally modulate mRNA stability and to coordinate transcription and decay ^13^. Furthermore, Rpb4-mediated post-transcriptional regulation also plays a major role in controlling the Environmental Stress Response (ESR) at both the transcription and mRNA decay levels ^13^. A recent study has demonstrated that disturbing Rpb4 dissociation from RNA pol II compromises the posttranscriptional roles of Rpb4 and leads to translation and mRNA decay defects ^17^.

Rpb4 acts as an RNA binding protein (RBP) by regulating gene expression from mRNA synthesis to mRNA degradation ^23^. Rpb4, and also Rpb7, are examples of RNA pol II components that act as RBPs ^7, 9, 17, 18, 23^. However, other RBPs associated with transcription machinery have been described, such as transcription elongation factors Spt4/5 ^23^, elements of the decaysome complex involved in mRNA degradation like Xrn1, Ccr4 or Dhh1 ^4, 8, 24^, or the Cbc elements of mRNA capping machinery ^25^. RBPs are elements that govern the post-transcriptional regulation of the mRNA life cycle in all living organisms that usually bind 3’ UTR elements of their targeted mRNAs, modulating their splicing, stability, translation and cellular location in eukaryotic organisms ^23, 26–33^. Different RBPs bind to specific and distinct sets of mRNAs that typically encode proteins with related biological functions or destined to similar subcellular localizations ^27, 32, 34^. Many RBPs shuttle between nucleus and cytoplasm ^32^. The distribution of RBPs between the nucleus and cytoplasm impacts gene expression through their interaction with their cytosolic or nuclear mRNAs ^32^.

Despite Rpb4–mRNA imprinting have been demonstrated ^7, 13^, very little is known about the specific mRNAs bound by Rpb4 and the molecular mechanisms governing mRNA life cycle regulation *via* Rpb4.

In this work, we investigated the precise role of Rpb4 in mRNA imprinting and degradation. The genome-wide analysis by RIP-Seq identified a set of almost 1500 mRNAs bound to Rpb4. We found that the average degradation rate of the Rpb4-bound mRNAs was higher than the overall one of mRNAs, which falls in line with a role for Rpb4 in mRNA stability. Notably a fraction of these transcripts are also targets of RPB Puf3. Puf3 is a well-studied example of RBPs of the PUF (Pumilio and FBF) family, a conserved family of mRNA regulators that bind to the 3’ UTR of many mRNAs to regulate mRNA stability and/or translation ^35–39^, which binds nuclear-encoded mitochondrial proteins ^40–42^. In line with this concerted Rpb4 and Puf3 association to a group of mRNA, our results demonstrated that Rpb4 and Puf3 show genetic and physical interactions. The integrity of the Puf3 protein depends on Rpb4. We also found that the Rpb4 and Puf3 association with mRNAs depends on each other and that Puf3 associates with chromatin in an Rpb4-dependent manner. Therefore, we conclude that Rpb4 and Puf3 cooperate to control their respective association with mRNAs and the mRNA stability of a specific group of mRNAs, which suggest a concerted role in mRNA imprinting. Our data also point out Rpb4 as a hub that cooperates with different RBPs in the mRNA synthesis and degradation interplay.

## MATERIALS AND METHODS

### Yeast strains, genetic manipulations, media and genetic analysis

Common yeast media, growth conditions and genetic techniques were used as described elsewhere ^43^.

Yeast strains and primers are listed in Supplementary Tables S1-S2, respectively.

### Isoform-specific RNA immunoprecipitation (isRIP)

RNA immunoprecipitation was carried out as described in ^44^. Briefly, two biological replicates of 1l cell cultures of the exponentially grown Rpb4-TAP-tagged strain (OD_600_ 0.8-0.9) were pelleted, washed twice with 25 ml of buffer A (20mM Tris-HCl, pH 8, 140 mM KCl, 1.8 mM MgCl_2_, 0.1% NP-40, 0.02 mg/ml heparin) and resuspended in 2 ml of buffer B (20 mM Tris-HCl, pH 8, 140 mM KCl, 1.8 mM MgCl_2_, 0.1% NP-40, 0.2 mg/ml heparin, 0.5 mM DTT, 1 mM PMSF, 50 U/μl RNasin Plus, 1X protease inhibitor cocktail [Complete; Roche]). Then cells were broken in a FastPrep24 agitator for 30 seconds at speed 5.5 and 2x for 20 seconds at 6.0, followed by a 1-minute break between each step by adding glass beads (425-600 μm, Sigma). Extracts were clarified by centrifuging twice at 7000 x g for 5 min. A 100 μl cell lysate aliquot was used as the INPUT control. Rpb4 was immunopurified from cell lysates by using 400 μl of Dyna M280 sheep anti-rabbit IgG beads (Invitrogen) for 2 h at 4°C with gentle rotation in a rotation mixer. After four 15-minute washes with buffer C (20 mM Tris-HCl, pH 8, 140 mM KCl, 1.8 mM MgCl_2_, 0.1% NP-40, 10% glycerol, 0.5 mM DTT, 1 mM PMSF, 50 U/μl RNasin Plus, 1X protease inhibitor cocktail [Complete; Roche]), RNA-Rpb4 complexes were eluted from beads by TEV protease cleavage of TAP-tag using 10 μl of 10 U/μl acTEV protease for 1 h at 21°C in a heatblock by shaking at 700 rpm.

The eluent was collected and RNA was isolated by extracting twice with one volume of phenol: chloroform: isoamyl alcohol (25:24:1), pH 4.7, and centrifuging at 13000 rpm for 1 min at room temperature. The upper phase was recovered and an equal volume of chloroform-isoamyl alcohol (24:1) pH 4.7 was added, gently mixed and centrifuged for 1 minute at 13000 rpm and room temperature. RNA was then precipitated from the upper phase with two volumes of 100% ethanol, 1/10 volume of NaAc 3M and 3μl of linear acrylamide as a carrier by incubating for 20 minutes on ice. Then RNA was pelleted by centrifugation at 14000 rpm for 20 minutes at 4°C and washed with 500 μl of 70% ethanol. After centrifugation at 13000 rpm for 1 min at room temperature, the pellet was dried and resuspended in 15 μl of milliQ H_2_O.

INPUT RNA extraction was performed with the RNeasy Mini Kit *(Quiagen)* according to the manufacture’s indications.

### 3T-fill sequencing and data analysis

Two biological replicates were used and the RNA from the total extracts (input) and immunoprecipitated samples (RIP) was sequenced. Constructions of sequencing libraries were carried out as previously described ^45^ from 1 μg RNA for inputs and 150 ng RNA for the RIP samples. Data were aligned to yeast reference genome version R64 for strain S288c using a custom computational pipeline, followed by the calculation of RIP enrichment using the DESeq2 R Bioconductor package as previously described ^44, 46^.

### mRNA extraction and reverse transcription

Total RNA from yeast cells was extracted and quantified as previously described from 50 ml of culture cells ^47^.

First-strand cDNA was synthesized using 1 μg of RNA with the iScript cDNA synthesis kit (Bio-Rad) following the manufacturer’s protocol. As a negative control for genomic DNA contamination, each sample was subjected to the same reaction without reverse transcriptase.

### Quantitative real-time PCR (RT-qPCR)

Real-time PCR was performed in a CFX-384 Real-Time PCR instrument (BioRad) with the EvaGreen detection system ‘‘SsoFast™ EvaGreen® Supermix” (BioRad). Reactions were performed in a total volume of 5 μl that contained the cDNA corresponding to 0.1 ng of total RNA. Each PCR reaction was performed at least 3 times with three independent biological replicates to gain a representative average. The 18S rRNA gene was used as a normalizer. Supplementary Table S2 lists the employed oligonucleotides.

### Isolation of the mRNA-associated proteins

mRNA crosslinking was carried out as described in ^13^ with some modifications. Briefly, 250 ml of cell cultures with an OD600 ~ 0.6-0.8 were exposed to 1200 mJ/cm^2^ of 254 nm UV in a UV crosslinker (Biolink Shortwave 254 nm), resuspended in 350 μl of lysis buffer (20 mM Tris, pH 7.5, 0.5 M NaCl, 1 mM EDTA, 1x protease inhibitor cocktail [Complete; Roche]) and 200 μl of glass beads (425-600 μm, Sigma), and were broken by vortexing for 15 min at 4°C. A lysate aliquot was used as the INPUT control. The lysate was incubated with 150 μl of oligo (dT)_25_ cellulose beads (New England BioLabs, cat no. S1408S) for 15 min at room temperature, washed 5 times with loading buffer (20 mM Tris–HCl, pH 7.5, 0.5M NaCl, 1mM EDTA) and once with low-salt buffer (10 mM Tris-HCl, pH 7.5, 0.1 M NaCl, 1 mM EDTA). The RNA-associated proteins were eluted by resuspending beads in 250 μl of elution buffer (20 mM Tris–HCl, pH 7.5) pre-warmed at 70°C by incubating 5 min at room temperature (x 2). The two eluate samples were mixed, lyophilized and resuspended in 20 μl of milliQ H_2_O. The mRNA-associated proteins were analyzed by SDS-PAGE and Western blot with the appropriate antibodies: anti-Rpb4 (Pol II RPB4 (2Y14); Biolegend), anti-Pgk1 (22C5D8; Invitrogen), anti-H3 (ab1791; Abcam); PAP (Sigm*a*).

### Protein extraction and TAP purification

Protein whole cell extracts and TAP purifications were performed as described in ^47^. Briefly, 150 ml of cells grown exponentially (OD_600_ 0.6-0.8) were pelleted and resuspended in 0.3 ml of lysis buffer (50 mM HEPES [pH 7.5], 120 mM NaCl, 1 mM EDTA, 0.3% 3-[(3-cholamidopropyl)-dimethylammonio]-1-propanesulfonate (CHAPS)), supplemented with 1X protease inhibitor cocktail (Complete; Roche), 0.5 mM phenylmethylsulonyl fluoride (PMSF), 2 mM sodium orthovanadate and 1 mM sodium fluoride. Cells were broken by vortexing (3 cycles, 5 min each) using 0.2 ml of glass beads (425-600 μm; Sigma)). TAP purification was carried out as previously described ^48^ with 2 mg of protein extract and 50 μl of Dynabeads Pan Mouse IgG (Invitrogen) per sample. The affinity-purified proteins were released from the beads by boiling for 10 min and performing western blotting with different antibodies.

For the RNase treatment, the cell-crude extracts (60 μg proteins) were incubated with 15 μg of RNase A, at 37°C for 1 h, before proceeding with the yChEFs method, as previously described ^49^.

### SDS-PAGE and western blot analyses

Protein electrophoresis and Western blot were carried out as described _in_ ^48^.

For the western blot analyses, the anti-Rpb4 (Pol II RPB4 (2Y14); Biolegend), anti-Pgk1 (22C5D8; Invitrogen), anti-H3 (ab1791; Abcam) and PAP *(Sigma)* antibodies were used.

### Yeast chromatin-enriched fractions preparation

The chromatin-enriched fractions preparation was carried out by the yChEFs procedure ^49, 50^ using 75 ml of exponentially grown YPD cultures (OD600 ~0.6–0.8). The final pellet obtained by this methodology was resuspended in 20 μl of 1x Tris-Glycine SDS sample buffer and incubated for 5 min at 100°C to obtain chromatin-bound proteins, which were analyzed by western blot with the different antibodies indicated above. The anti-histone H2A (39235, Active Motif) and anti-phosphoglycerate kinase Pgk1 (459,250; Invitrogen) antibodies were employed to detect H3 histone, used as a positive control, and Pgk1 as a negative control of cytoplasmic contamination.

### Chromatin immunoprecipitation and sequencing (ChIP-Seq)

Chromatin immunoprecipitation was performed using three biological replicates for strains Puf3-TAP WT and Puf3-TAP *rpb4Δ*, and two for the WT strain. For each sample, 100 ml of cells were grown in YPD medium to OD600 ~ 0.7. The SNF21-TAP-natR *S. pombe* strain (spike-in control, SP641, *h90 ura-r TetR-tup11 KanR-tetO-snf21-TAP-natR ura4-D18)* was grown in YES medium to OD_600_ ~ 0.7. Cells were crosslinked by adding 1% formaldehyde (Thermo Fisher) directly to the culture and shaking for 15 minutes. Formaldehyde was quenched for 5 min by adding 0.125 M glycine. Cells were pelleted, washed 4 times with cold TBS and flash-frozen.

Pellets were thawed on ice, resuspended in 500 μl of zymolyase solution (1.2 M sorbitol, 10 mM Tris-HCl, pH 8, 10 mM CaCl2, 1% v/v betamercaptoethanol, 0.1 U/μl zymolyase (Zymo Research)) per pellet and incubated at 37°C for 30 minutes with shaking at 500 rpm to digest cell walls. Spheroplasts were isolated by centrifugation (5 min at 3500xg) and. resuspended in 540 μl of lysis buffer (50 mM HEPES-KOH, pH 7.5, 140 mM NaCl, 1 mM EDTA, 1% v/v Triton X-100, 0.1% w/v sodium deoxycholate, 1 mM PMSF, 1 mM benzamidine, 1:100 yeast protease inhibitor cocktail (Sigma Aldrich, P8215)). Lysates were sonicated in a Covaris ME220 instrument in 130 μl screw-cap microtubes for 3 min to 100-500 bp (PIP = 75, DF = 15%, 1000 cycles per burst, 9°C). The sonicated lysates were centrifuged at 17000xg for 10 minutes to pellet the insoluble material. For IP, an equal amount of chromatin from each sample was taken based on the estimated DNA concentration, and *S. pombe* sonicated chromatin was spiked to approximately 1% in relation to that amount (*S. pombe* chromatin was obtained as described above, except for using a 2-fold volume of zymolyase solution for twice the time).

Next 250 μl of Dynabeads Pan Mouse IgG (Thermo Fisher) per sample were washed 3 times with PBS/BSA (PBS with 5 mg/ml BSA (Sigma Aldrich)), resuspended in the sonicated chromatin solution and rotated overnight at +4°C. They were washed 3 times with lysis buffer, lysis buffer containing 500 mM NaCl and LiCl wash buffer (10 mM Tris–HCl, pH 8, 1 mM EDTA, 250 mM LiCl, 0.5% NP-40, 0.5% sodium deoxycholate), and once with elution buffer (50 mM Tris-HCl, pH 8, 1% SDS, 10 mM EDTA). Chromatin was eluted from beads in elution buffer at 65°C with shaking for 20 minutes. The eluted material was diluted 5-fold to lower the SDS concentration to 0.2%, then 1:100 RNase cocktail (Thermo Fisher, AM2286) was added and samples were incubated at 37°C for 45 minutes to digest RNA. SDS up to a total of 0.5% and Proteinase K up to 0.5 mg/ml (Thermo Fisher, AM2546) were added, and samples were shaken overnight at 65°C to reverse the crosslinked DNA. Input controls (10 μl sonicated chromatin) were subjected to RNA digestion and crosslinking reversion in the same way. DNA was purified with the QIAquick Gel Extraction kit (QIAGEN) and eluted in 50 μl of water.

Sequencing libraries were prepared using the NEBNext Ultra™ II DNA Library Prep Kit (NEB). Four PCR cycles were used for the input samples and 17 cycles for the IP samples. The resulting libraries were pooled and dual size-selected with 0.6X and 1.2X AMPure XP (Beckman Coulter) bead ratios. The pool was sequenced in NextSeq 500 with paired-end with a read length of 38 bases.

Fastaq files were mapped to a combined *S. cerevisiae* (R64) and *S. pombe* genome using bowtie2 with default settings. The mapped files were deduplicated with Picard and filtered by mapping quality (MAPQ>2). Peaks were called with MACS2 (--nomodel mode, --extsize 170, −q 0.05) using all the Puf3-TAP and Puf3-TAP *Δrpb4* samples by taking the WT control as input. Metagene profiles were plotted with VAP ^51^ over all the *S. cerevisiae* genes and the genes whose mRNA was bound by Puf3.

## RESULTS

### Rpb4 physically interacts with mRNAs

To investigate the association of the RNA pol II specific subunit Rpb4 with mRNAs *in vivo*, we performed RIP experiments using the previously described isRIP methodology (isoform-specific RNA immunoprecipitation) ^44^, which allows the quantification of mRNA isoforms that were immunoprecipitated with a RBP.

For that purpose we used an Rpb4 TAP-tagged yeast strain grown in YPD medium at 30°C, and immunoprecipitated the RNA natively bound to Rpb4 without cross-linking procedure. Two biological replicates were used and the RNA from total extracts and the Rpb4-immunoprecipitated samples were sequenced by genome-wide 3’ isoforms sequencing ^45^, measuring the enrichment of each mRNA in the immunoprecipitated (IP) fraction relative to the input sample. As expected, the biological replicates showed high reproducibility (Fig. S1). By using the “DEseq2” software ^52^, the normalized fold change in the IP samples was calculated (Fig.1A). To define the Rpb4-bound transcripts, a false discovery rate (FDR) of < 10% was applied, as previously described ^44^, along with a specific cut-off consisting of Log2 fold-change of RIP vs. input higher than 0 and padj<0.1. We identified 1458 mRNAs differentially enriched in the RIP samples over the total extract. Notably, when binding stringency was increased to a 1.5 fold change (padj<0.1), enrichment persisted and then 777 Rpb4-bound mRNAs were identified. However, as Rpb4 was expected to bind a significant fraction of the transcriptome (unlike other RPB), we decided to adopt the more permissive criteria.

**Figure 1:**
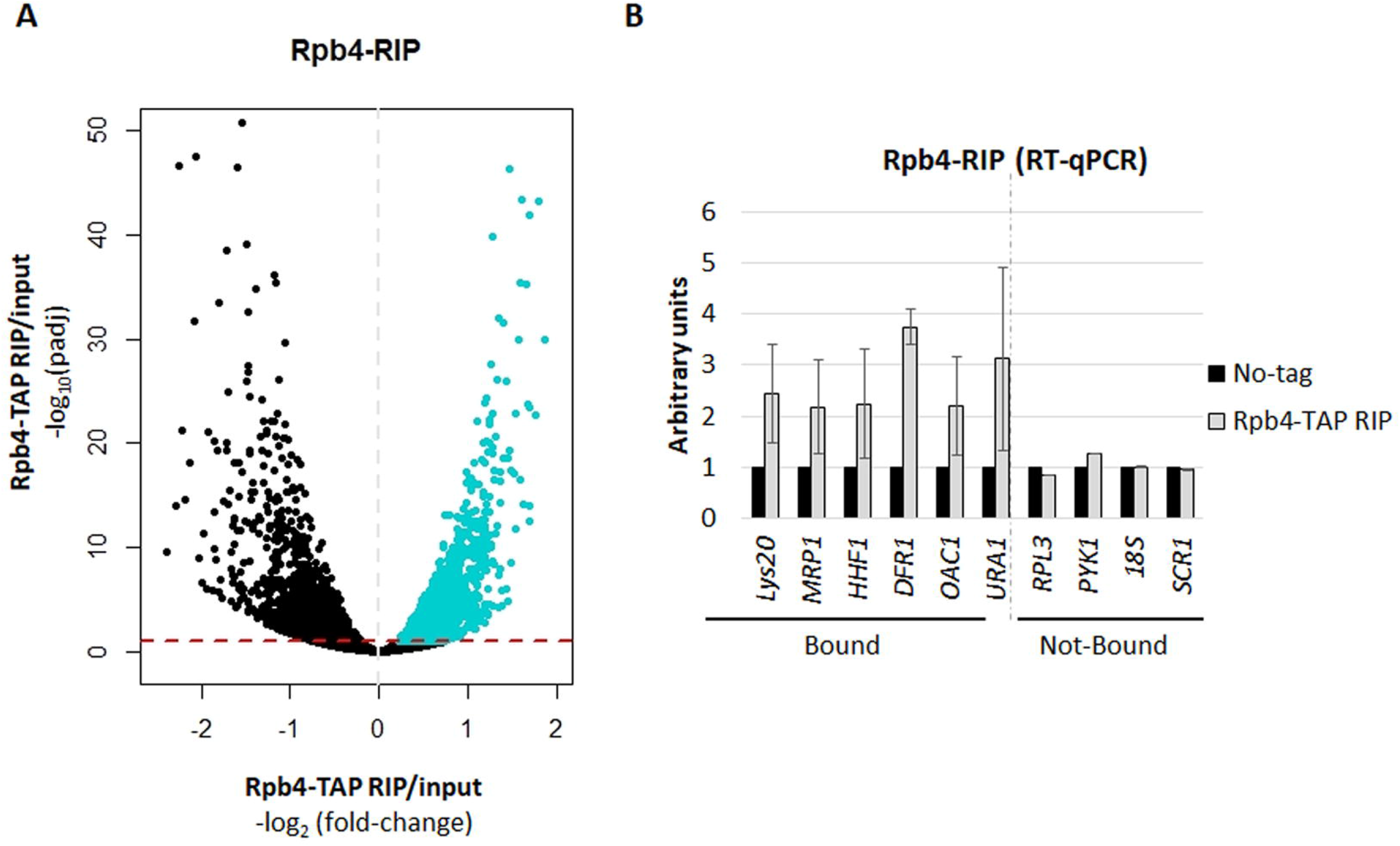
RNA immunoprecipitation (RIP) for Rpb4. **A)** Volcano-plot generated to identify the significantly bound mRNAs by Rpb4 (blue), calculated by the DESeq2 software (RIP vs. input) ^52^. A specific cut-off was applied to define the Rpb4-bounded transcripts consisting of Log_2_ Fold-Change of RIP vs. input higher than 0 (dashed gray line) and padj<0.1 (dashed red line). **B)** RT-qPCR to analyze several Rpb4-bound and unbound mRNAs from the RIP experiments corresponding to the Rpb4-TAP and the no-TAP tag control strains. Data are shown as the average and standard deviation (SD) of at least three independent experiments (RIP vs. input). rRNA 18S was used as a normalizer.

Furthermore, the RIP-Seq experiment results were corroborated by analyzing the enrichment of some of the identified Rpb4-bound mRNAs, by RT-qPCR, in relation to both a no TAP-tag control strain and to some Rpb4 non-bound mRNAs, in the input and immunoprecipitated samples. As shown in Figure 1B, all the analyzed Rpb4-bound mRNAs were enriched in relation to the no TAPtag control strain. As expected, the analysis of some Rpb4 non-bound mRNAs showed no enrichment either for the Rpb4 TAP-tagged yeast strain or the no TAP-tag control strain (Fig. 1B). In addition, the highly abundant *18S* and *SCR1* transcripts of RNA pol I and III, respectively, showed no enhanced signal in the RIP samples, which confirmed that Rpb4 was bound to specific RNA pol II transcripts, as expected from an RNA pol II specific subunit.

We performed a functional analysis of GO categories of the biological process for the Rpb4-bound mRNA identified in our genome-wide analysis, by using the String software ^53^. We found, statistically significant, categories corresponding to mitochondrion, such as mitochondrial translation, mitochondrial gene expression or mitochondrial transport (Table S3). In addition, other statistically significant categories related to transmembrane transport and metabolic processes were detected.

### Rpb4 and Puf3 associate with a common group of target mRNAs

As shown above, the Rpb4-bound mRNAs are enriched for the mitochondrion-related GO categories. In addition, the mRNA encoding mitochondrial proteins have been described to be bound by the member of the Pumilio and FBF family RBPs Puf3 at their 3’UTR region ^40–42, 44^.

These results suggest a potential relation between Rpb4 and Puf3. Accordingly, we speculate that Rpb4-bound mRNAs could also be imprinted by Puf3. To explore it more in detail we compared our RIP data to different available datasets of Puf3 mRNA interactors identified by distinct methodologies. We firstly compared our data by the RIP-Seq procedure to data from two different Puf3 RIP-Seq analyses ^44, 54^. As shown in Figure 2A and Figure S2, the Rpb4-bound transcripts significantly matched the Puf3-bound mRNAs compared to the dataset defined by Kershaw et al ^54^, and to a lesser extent compared to the more stringent dataset defined by Gupta et al ^44^, probably due to few identified Puf3-bound mRNAs as a consequence of the high fold-change used. To gain more insight, we also compared our data to other datasets of Puf3-bound transcripts identified by other methodologies, HITS-CLIP ^55^, RIP-ChIP ^56^ and PAR-CLIP ^57^. As shown in Figure 2B and Figure S2, again, a significant overlap was detected. To be more stringent, we also performed the same kind of analysis by selecting a 1.5 fold-change for our RIP data (RIP vs. input). In this case, the data from our analysis significantly matched the other used datasets, with similar or even higher percentage of overlapping (Figure S2).

**Figure 2:**
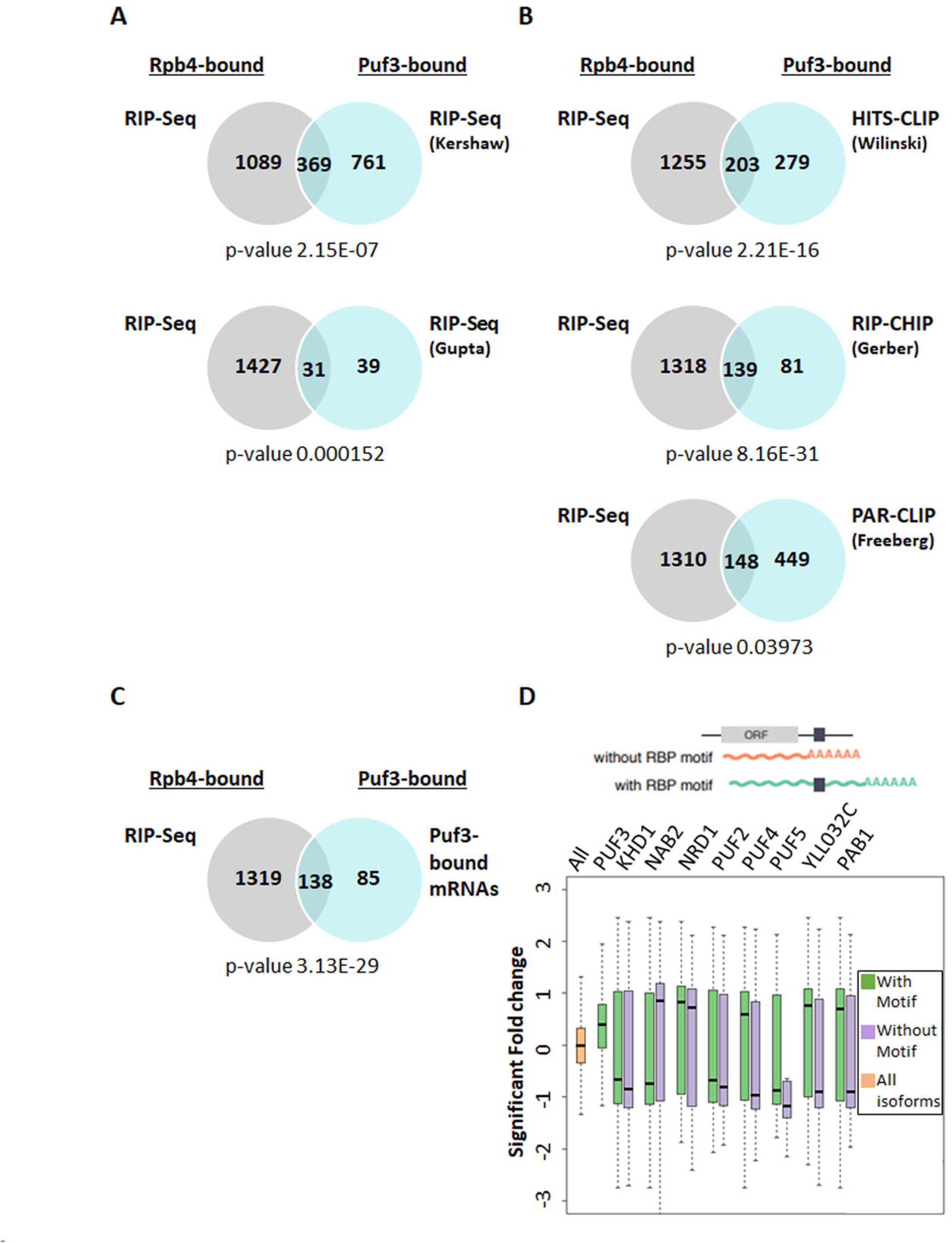
Rpb4-bound mRNAs correlates with Puf3-interactors. **A)** Venn diagrams showing the overlapping found between the mRNAs bound by Rpb4 and Puf3 identified by RIP-Seq ^44, 54^. **B)** Venn diagrams showing the coincidence between the Rpb4-RIP and Puf3-bound mRNAs identified by HITS-CLIP ^55^, RIP-CHIP ^56^ and PAR-CLIP ^57^. **C)** Venn diagrams depicting the common mRNAs bound by Rpb4 and Puf3 (for Puf3, they were selected as those in common in at least 3 of the 5 datasets indicated in B). **D)** Boxplots showing the distribution of significant Rpb4 Log2 enrichment over the two different categories of polyadenylation isoforms belonging to genes with a particular predicted RBP binding motif (schematic on top). In the boxplots, all the polyadenylation isoforms are depicted in orange, the polyadenylation isoforms of the genes containing the predicted motif of the specified RBP are denoted in green while all the other polyadenylation isoforms of the same genes are shown in purple. The isoform-specific association of Puf3 with mRNAs has been previously defined ^44^ and the other RBP motifs were taken from ^70^.

To gain more insight into these analyses, we then defined as Puf3-bound mRNAs all the Puf3 targets that were common in at least three of the five datasets used, which are presented in Figure 2 ^44, 54–57^. Accordingly, we defined 223 mRNAs as Puf3-bound mRNAs (Table S4). Notably, most of Puf3-bound mRNAs, 138 (62 %), matched the Rpb4-bound mRNAs identified by RIP-Seq in our analysis (Figure 2C). Notably when a more stringent analysis was applied to identify the Rpb4-bound mRNAs (Log_2_ fold-change of RIP vs. input above 0.58 and padj<0.1), 104 (47%) of the Rpb4-bound mRNAs matched the Puf3-bound mRNAs. As expected, and in line with our above results and with previous results for Puf3 ^40–42, 44^, the analysis of GO categories of Biological process, by using String software ^53^, of the Rpb4 and Puf3 common-bound mRNAs showed that the Biological process was related mainly to mitochondria and mitochondrial translation (Supplementary Table S5).

We also analyzed whether the Rpb4-bound transcripts were enriched in mRNAs with canonical Puf3 binding motif. In addition, and taking that Rpb4 imprints almost 1500 mRNAs (only 138 in common with Puf3), we also extended this analysis to different RBPs. Furthermore, we hypothesized that if an RBP and Rpb4 interaction existed, this would reflect a significant change in the isoform-specific RIP enrichment. To test this hypothesis, we compared our Rpb4-TAP RIP experiment data at the 3’ mRNA Isoforms level by separating the isoforms based on either their direct interaction with an RBP, such as Puf3 ^44^, or the presence of an RBP motif indicative of a putative interaction ^58^. We found that the 3’ isoforms which interacted with Puf3, Puf4, Puf5, Khd1, Nrd1, Pab1 and YLL032C had a significantly enhanced RIP signal (FDR <0.05), while those that interacted with Nab2 had a significantly reduced RIP signal (FDR <0.05) (Figure 2D). Notably, our results demonstrated that the Rpb4-bound mRNAs clearly correlated with the presence of the Puf3 binding motifs (Figure 2D), but also, likely, with other RBPs. This demonstrates that an interaction with specific RBPs is associated with the Rpb4 RIP signal possibly through the imprinting mechanism.

Taken together, the above data indicate a functional relation between Rpb4 and Puf3 in the imprinting of at least a set of common mRNA targets, which is indicative of a coordinated role for these RPBs during this process.

### Rpb4-bound mRNAs decay faster

The cells lacking Rpb4 or *rpb1* mutants *(rpo21-4 and rpb1-84)* that affect the Rpb4 association with RNA pol II reduce Rpb4-mRNA imprinting and generally lead to increased mRNA stability. This demonstrates the role of Rpb4 in regulating mRNA stability ^7, 13, 14^, which probably depends on the association between Rpb4 and general mRNA decay machinery Pat1/Lsm1-7 ^14^.

Hence we wondered if a correlation between mRNA stability and its association with Rpb4 could exist. To explore this possibility, we compared our data from the Rpb4-TAP RIP experiments with previously published decay rate data obtained by MIST-Seq (Measurement of Isoform-Specific Turnover using Sequencing), which showed the half-life of polyadenylation or 3’ transcript isoform in yeast ^44^. Notably as shown in Figure 3A, the 3’ transcript isoforms that were enriched in the Rpb4 RIP experiments had lower half-lives than the overall mRNA. Consequently, these data suggest that the Rpb4-bound mRNAs degrade faster than global mRNA, which falls in line with previous data showing that decreasing mRNA imprinting by Rpb4 provokes increased mRNA stability ^10, 13^.

**Figure 3:**
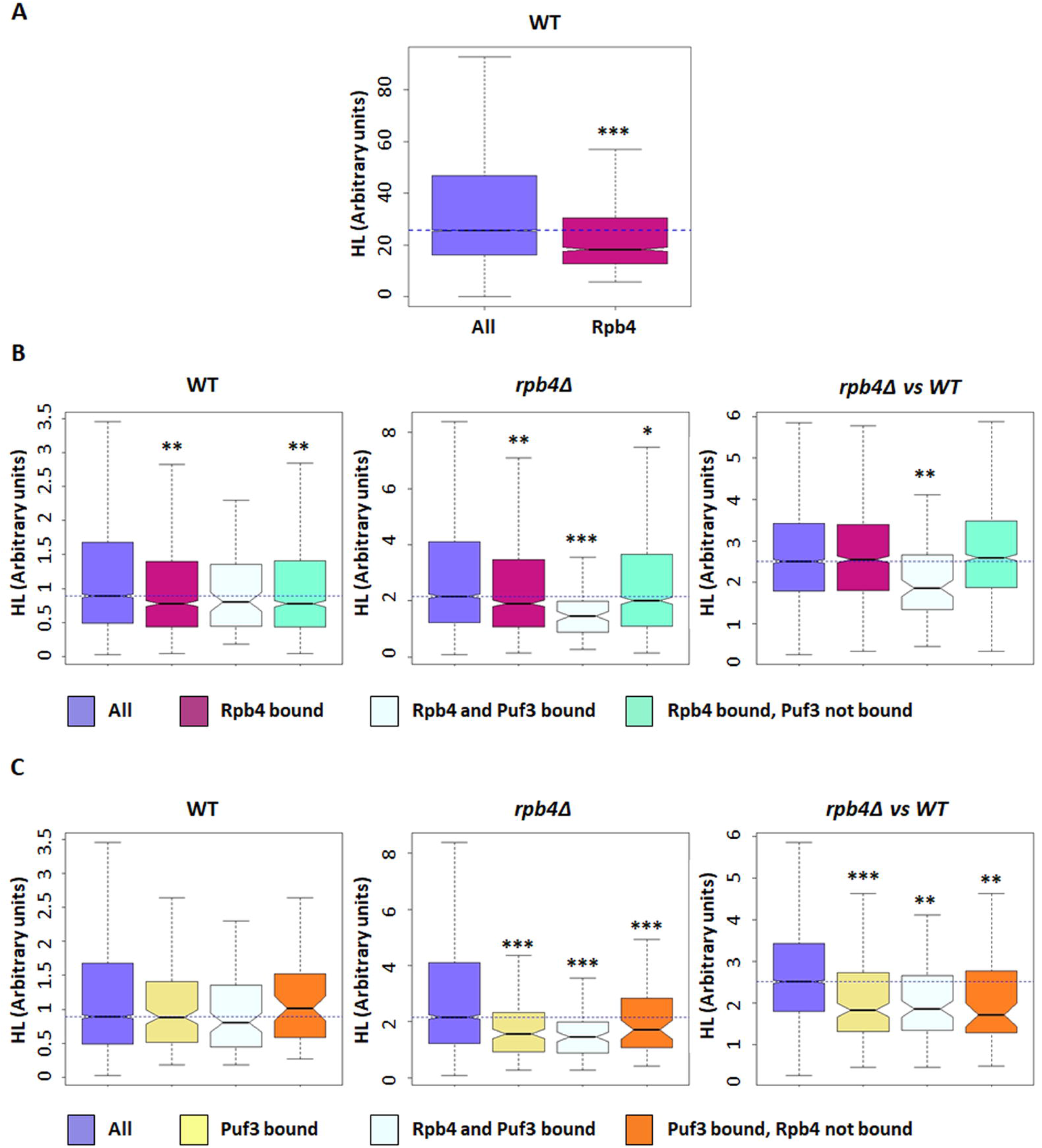
Rpb4-bound mRNA show a higher decay rate. **A)** Boxplots showing the half-lives of the general polyadenylation isoforms of a wild-type strain ^44^ for the Rpb4-bound mRNAs. **B)** Boxplot depicting the half-lives of the Rpb4-bound mRNAs in a wild-type strain (left panel), the *rpb4Δ* mutant strain (middle panel), or the *rpb4*ZVwt ratio (right panel). **C)** Boxplot showing the halflives for the Puf3-bound mRNAs in a wild-type strain (left panel), the *rpb4Δ* mutant strain (middle panel) or the *rpb4Δ*/wt ratio (right panel). Datasets used in B and C correspond to ^13^.* Wilcoxon-test p-value < 10^−3^; ** p-value < 10^−5^; *** p-value < 10^−9^. Each group of analyzed mRNA was compared to the group “all”.

According to these results, we speculated that Rpb4-bound and Rpb4/Puf3 commonly-bound mRNAs could be differentially affected by lack of Rpb4 at the degradation/stability level in relation to the overall cellular mRNA. To test this hypothesis, we used the published wild-type and *rpb4Δ* mRNA half-life dataset, from the cells grown in YPD at the permissive temperature of 30°C ^13^, obtained by the GRO method ^59^. As seen in Figure 3B (left panel), corroborating the data in Figure 3A, the Rpb4-bound mRNAs showed lower half-lives than the overall mRNA in the wild-type strain at the permissive temperature of 30°C.

However, Rpb4/Puf3 commonly-bound mRNAs did not show significant differences in mRNA stability with respect to the overall mRNAs, while it was the case for Rpb4 (non Puf3)-bound mRNAs (Figure 3B, left panel). When we analyzed the mRNA half-lives in the *rpb4Δ* mutant strain, a significant general increase in mRNA stability was observed (Figure 3B, middle panel) in agreement with previous reported data ^13^. Strikingly, this increase was significantly less marked for the Rpb4-bound mRNAs that were also bound by Puf3 (Figure 3B, middle panel). The less marked stability of the Rpb4-bound mRNAs *versus* the whole population was, however, identical when comparing the cells with (wt) or without (*rpb4*Δ) this protein, except for the Rpb4/Puf3 commonly-bound mRNAs (Figure 3B, right panel). These results indicate that Rpb4 has no specific effect on the stability of its mRNA targets, but a global one. However, lack of Rpb4 has specific effect on the targets shared with Puf3. It is worth noting that Puf3 mRNA targets were destabilized in the cells grown exponentially in YPD medium ^37, 38^. Thus we concluded that the Puf3-mediated decay of its targets is enhanced in the absence of Rpb4.

As a set of the Puf3 targets (as previously define) were not bound by Rpb4, we also explored the half-lives of the Puf3-bound mRNAs from the *rpb4Δ* mutant and its isogenic wild-type strain, using the same datasets. Unlike the Rpb4-bound mRNAs, the Puf3-bound mRNAs showed no significant differences in the half-lives in relation to the overall mRNAs in the wild-type strain (Figure 3C, left panel). In addition, all the Puf3 targets (both those targeted and not by Rpb4) displayed a similar less marked increase in their stability than the overall mRNA population when Rpb4 lacks (Fig. 3C, middle and right panels). This finding indicated that the effect of Rpb4 on the Puf3 targets does not require the simultaneous binding of both proteins to mRNAs.

### Rpb4 physically and genetically interacts with Puf3

The above data and the demonstrated role of Rpb4 and Puf3 in mRNA stability ^7, 13, 38^ suggest a coordinated role of Rpb4 and Puf3 in mRNA imprinting.

To test this hypothesis, we first investigated whether Rpb4 and Puf3 physically interacted. To do so, we performed TAP pulldown from a strain containing a fully functional Puf3-TAP tagged protein and analyzed the Rpb4 association using specific anti-Rpb4 antibodies. As shown in Figure 4A, Rpb4 interacted with Puf3-TAP, although this interaction seemed weak or transitory compared to the positive control employed in the experiment. As a positive control in the experiment, a clear interaction between Rpb4 and Rpb3 (subunit of RNA pol II) was observed when performing Rpb3-TAP tagged protein pulldown. Conversely, and as a negative control, neither non-specific binding between Rpb4 and Imd2-TAP (inosine monophosphate dehydrogenase that catalyzes the rate-limiting step in GTP biosynthesis ^60^) nor non-specific adsorption to the resin (no tag control) took place. We also investigated the association of Rpb4 with other RBPs by pulldown with the TAP-tagged versions of different RBPs (Figure 4A). The results indicate that Rpb4 interacted with Puf2 and Nsr1, albeit to a lesser extent than with Puf3. Conversely, no clear interaction among Rpb4 and Pub1, Puf4, Puf5, Nrd1, and Vts1 was observed. Furthermore, in agreement with the Rpb4 and Puf3 physical interaction, a negative genetic interaction between mutants *rpb4Δ* and *puf3Δ* was observed as the *puf3Δ* mutation aggravated the slow growth phenotype of the single *rpb4Δ* mutant (Fig. 4B).

**Figure 4:**
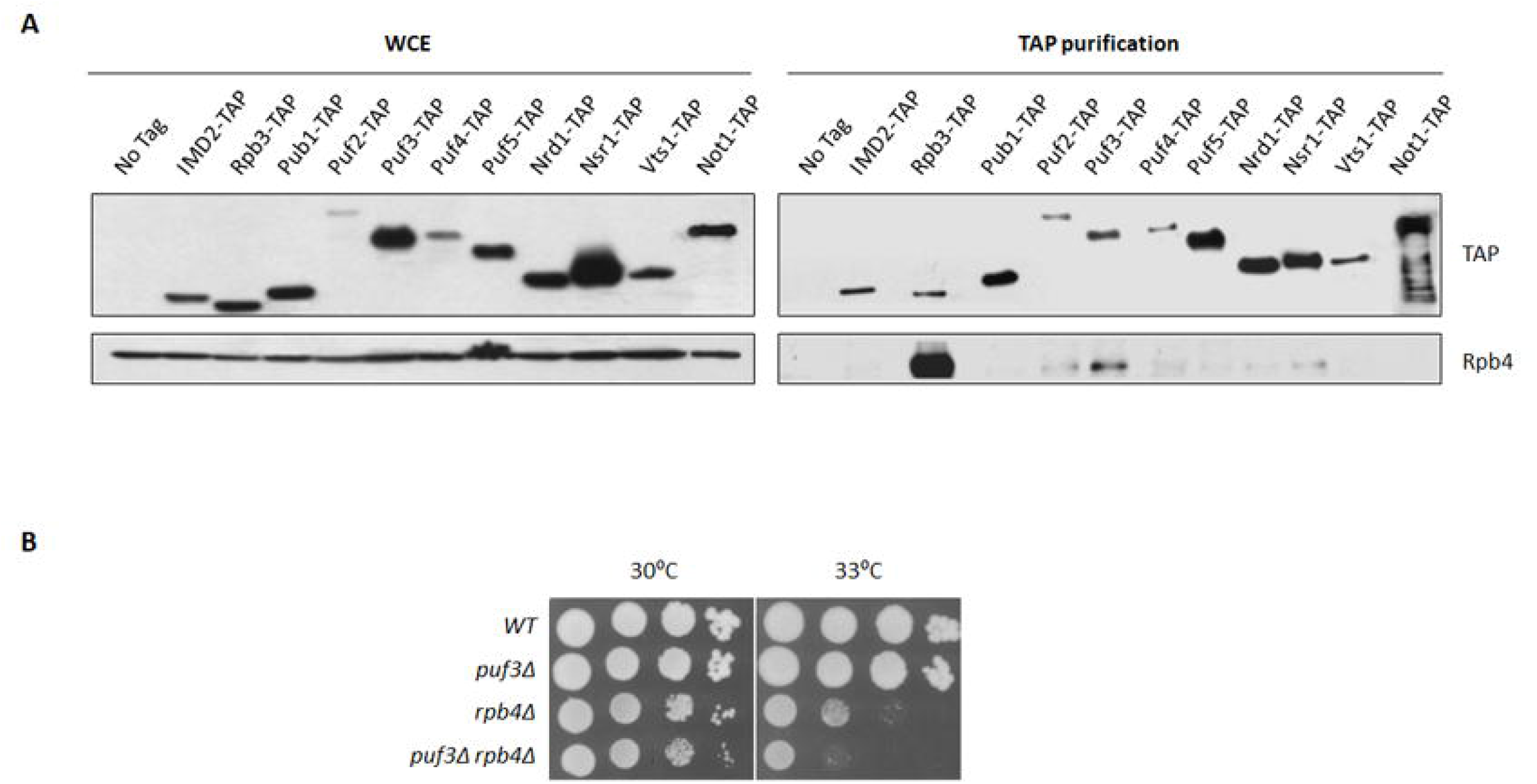
Rpb4 physically and genetically interacts with Puf3. **A).** Wholecell extract and TAP pulldown of the different RBPs TAP-tagged strains grown in YPD medium at 30°C. The resulting pulldowns were analyzed by western blot with antibodies against Rpb4 and TAP-tag. No tag strain was used as a negative control and the Imd2-TAP strain was employed as a control of nonspecific binding. The Rpb3-TAP strain was tested as a positive control for Rpb4 binding. **B)** Growth assay of the wild-type and *puf3Δ* mutant strains in combination with *RPB4* deletion in YPD medium at the indicated temperatures.

All these data collectively indicate a functional association between Rpb4 and Puf3 that could mediate a common imprinting mRNA mechanism, and suggest that this could also account for other RBPs. Finally, as the Rpb4 and Puf3 interaction was the strongest between Rpb4 and RPBs, we decided to focus on dissecting this interaction.

### Puf3 integrity depends on Rpb4

The above data prompted us to investigate the functional relation governing Rpb4 and Puf3 interaction. To explore it, we analyzed whether lack of Rpb4 could influence the Puf3 protein level. Protein Puf3-TAP was analyzed by western blot in crude extracts of *rpb4Δ* and its isogenic wild-type strains that contained a fully functional Puf3-TAP tagged protein, and were also grown in YPD at 30°C. Strikingly, a much faster migrating band (~100 KDa) appeared in the *rpb4Δ* strain (Fig. 5A), while the full-length Puf3-TAP band (~120 KDa) decreased.

**Figure 5:**
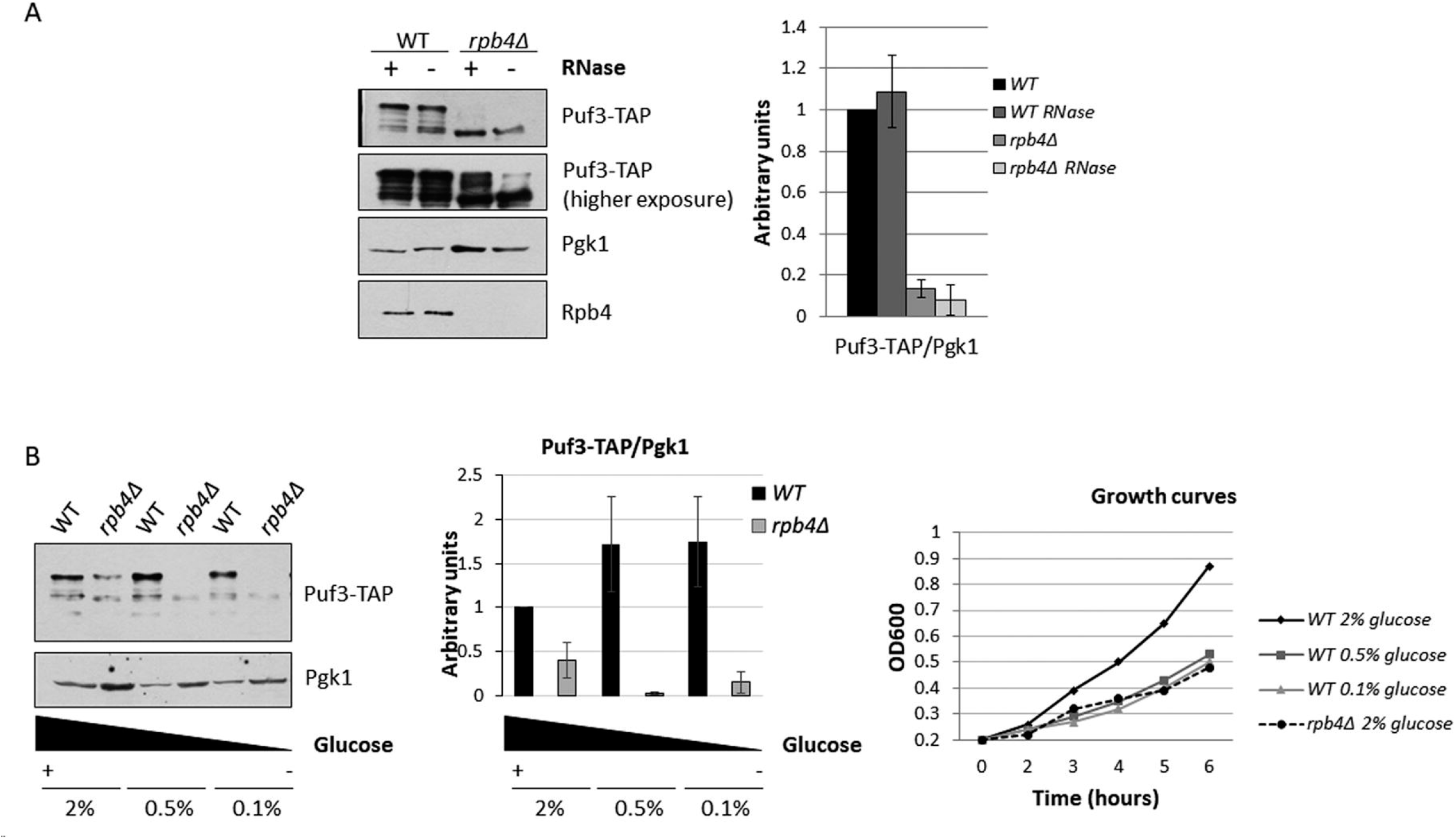
Puf3 integrity and *PUF3* mRNA levels depend on Rpb4. **A)** Wholecell extract of the Puf3-TAP tagged wild-type and *rpb4Δ* mutant strains with and without RNase treatment for 1 h at 37°C. Extracts were analyzed by western blot with antibodies against Rpb4, TAP-tag and Pgk1 used as a control. **B)** Western blot showing the Puf3-TAP and Rpb4 proteins in whole-cell extracts of the Puf3-TAP tagged wild-type and *rpb4Δ* mutant strains grown in YPD medium at 30°C at different glucose concentrations (left panel). Growth curves of the wild-type and *rpb4Δ* mutant strains in YPD medium containing the indicated glucose concentrations at 30°C.

It has been previously demonstrated that Puf3 is phosphorylated upon glucose depletion to modulate the fate of its target mRNAs from degradation to translation ^37^. However, the faster migrating form of Puf3 that we observed did not seem to account for differences in Puf3 phosphorylation as no differences in Puf3 migration were observed upon glucose depletion (see below and Fig. 5B). Moreover, as the TAP tag of Puf3, which is recognized by the anti-TAP antibody, is located at the C-terminal part of the protein, the faster migrating Puf3 band should correspond to the proteolysis of the N-terminal domain, and would probably correspond to part of the Puf3 domain phosphorylated upon glucose depletion ^37^. Nevertheless, the faster migration band did not seem to result from only a difference in Puf3 phosphorylation, because incubating the cell crude extracts of the wild-type cells (grown at 30°C in YPD) with phosphatase led to neither changes in the amount of the full-length Puf3-TAP band (~120 KDa) nor in the faster migrating band appearing (not shown).

In addition, and in order to rule out the notion that the appearance of the faster migrating band could be caused by the expression of a cryptic transcript leading to the production of a truncated protein ^61^, we confirmed by RT-qPCR that the production of full length transcript of *PUF3* was similar in the WT and *rpb4Δ* strain (Supplemental Figure S3).

We also analyzed whether the overall amount or Puf3 integrity depended on its association with RNA. To do so, we treated cell extracts with RNAse for 1 h and analyzed tagged Puf3-TAP by western blot. As shown in Figure 5A, RNAse treatment did not significantly affect the Puf3 levels in a wild-type strain. Similarly, the amount of the faster migrating Puf3 band observed when Rpb4 was lacking did not seem to be significantly altered.

The *rpb4Δ* mutant shows a slow growth phenotype ^13, 21^. In order to rule out whether the Puf3 faster migrating band could result from an indirect effect due to the slow growth of the *rpb4Δ* mutant cells, we analyzed by western blot the Puf3-TAP protein in a wild-type and *rpb4Δ* strain growing at different glucose concentrations as the carbon source to modulate their doubling time (Figure 5B). At 0.5% or 0.1% glucose, the wild-type strain showed a slow growth phenotype comparable to that of the *rpb4Δ* mutant at 2% glucose (Figure 5B, right panel). However as we can see in Figure 5B (left panel), increasing doubling time did not significantly alter Puf3 levels, nor the faster migrating Puf3 band appearing in a wild-type strain, unlike that observed upon *RPB4* deletion (Fig. 5B, left panel).

Taken together, lack of Rpb4 led to a faster migrating Puf3 band of about 100 KDa appearing, which suggests Puf3 proteolysis.

### Puf3 integrity depends on Rpb4 associated with chromatin-bound RNA pol II and not on Rpb4-mRNA imprinting

The above data suggest that Rpb4 is necessary for Puf3 integrity. Then, in order to investigate whether the alteration in Puf3 migration could depend on either the Rpb4 subunit bound to chromatin-associated RNA pol II or free Rpb4, we took advantage of the a *rpb1-84* mutant strain, which affects RNA pol II assembly and partially dissociates Rpb4 from the rest of the enzyme ^48^. This mutation leads to different chromatin-associated RNA pol II subcomplexes, some of which lack Rpb4, without altering Rpb4 cellular content ^48^. Then the Puf3-TAP protein was analyzed by western blot in crude extracts of the *rpb1-84* mutant strain and its isogenic wild-type strain containing this TAP-tagged protein. As shown in Figure 6A, the Puf3-TAP faster migrating band (100 kDa) was also observed in the *rpb1-84* mutant strain. These results indicate that the appearance of the faster migrating Puf3 band depends on the Rpb4 association with chromatin-bound RNA pol II. Furthermore, bearing in mind that the *rpb1-84* mutant and its wild-type strains have different genetic backgrounds than previously shown (above experiments), these data indicate that this phenomenon is independent of strain’s genetic background.

**Figure 6:**
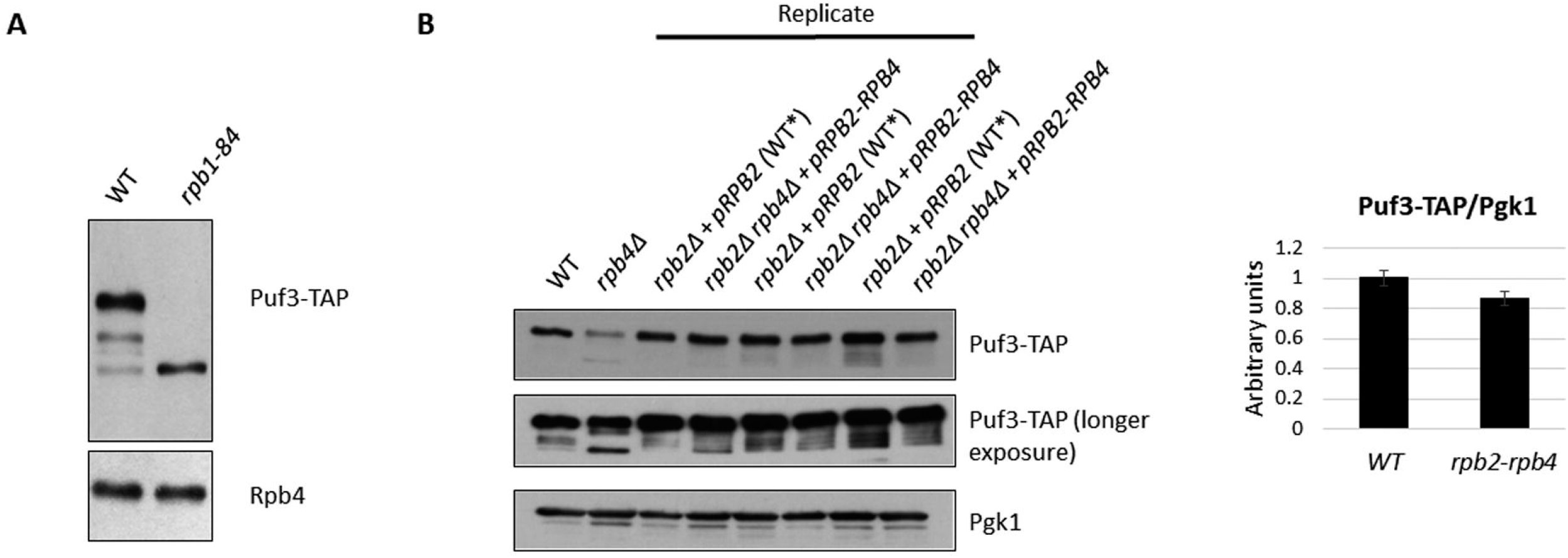
Puf3 integrity is compromised by lack of Rpb4. Whole-cell extract of the Puf3-TAP tagged wild-type and *rpb1-84* mutant strains analyzed by western blot with antibodies against Rpb4 and TAP-tag. **B)** Whole-cell extract of the Puf3-TAP tagged *rpb2Δ* strain expressing *RPB2* from a plasmid (WT*), and the Puf3-TAP tagged *rpb2Δ rpb4Δ* strain expressing an *RPB2-RPB4* fusion gene from the same plasmid analyzed by western blot with antibodies against TAP-tag and Pgk1 as a control (left panel). An *rpb4Δ* mutant strain and its wild-type isogenic strain (WT) were used as a control. Protein quantification by gel densitometry (right panel).

It has been demonstrated that Rpb4 fused to Rpb2 associates with RNA pol II and overcomes the transcriptional defect of the *rpb4Δ* mutant strain, ^17, 21^, although only partially recovers its effect on the mRNA degradation rate ^17^. Moreover, in the strain that expresses the Rpb2-Rpb4 fusion protein, a small fraction of Rpb2-Rpb4, but also of free Rpb4, are able to imprint mRNAs ^17^. So by using this construction, we also explored whether Rpb4-dependent mRNA imprinting would be necessary for Puf3 integrity. To do so, we used two Puf3-TAP tagged stains, the *rpb2Δ rpb4Δ* strain expressing a Rpb2-Rpb4 fusion protein from a plasmid ^17, 21^ and the isogenic *rpb2Δ* strain expressing the Rpb2 protein from the same plasmid and Rpb4 from the chromosomic *RPB4* gene, considered to be wild type. As shown in Figure 6B, when cells expressed the Rpb2-Rpb4 fusion protein, no faster migrating Puf3 band was observed in whole cell crude extracts, which indicates that Rpb2-Rpb4 suffices to maintain Puf3 integrity. Similar results were observed in the isogenic cells, indicated above, considered to be wild type (Figure 6B, WT*). As a control, an *rpb4Δ* strain showed the faster migrating band (~100 kDa), which was not observed in its isogenic wild-type strain (WT). By taking these data and those for the *rpb1-84* mutant, we speculate that mRNA imprinting by Rpb4 is not necessary for maintaining Puf3 integrity.

Rpb4 has been proposed to interact with mRNA in the context of RNA pol II during transcription ^7, 10, 13^. In order to decipher whether lack of Puf3 could alter the Rpb4 association with chromatin, we purified chromatin-enriched fractions by the yChEFs procedure ^49, 50^ and analyzed the Rpb4 association by western blot. Our results indicated that Rpb4 binding to chromatin did not seem dependent on Puf3 (Figure 7A), as evidenced by similar Rpb4 levels associated with chromatin in the *puf3Δ* and wild-type strains. By employing specific antibodies against 3-phosphoglycerate kinase (Pgk1) as a control of cytoplasmic protein, and against histone-3 as a control of chromatin-associated proteins, we observed that chromatin was successfully isolated, and similar chromatin levels were purified and analyzed in the wild-type and mutant cells.

**Figure 7:**
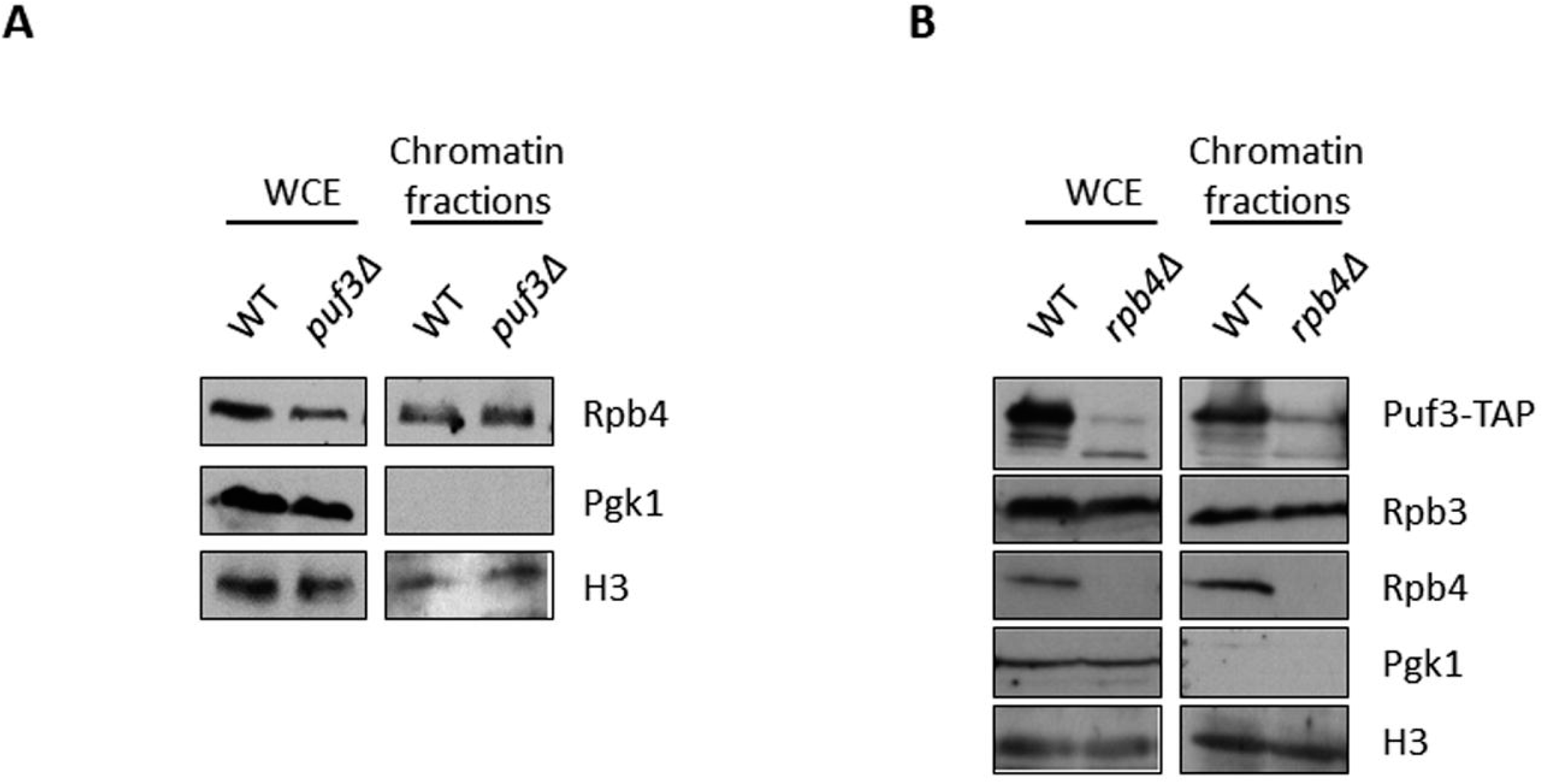
Puf3 and Rpb4 association with chromatin. **A)** The whole-cell extract and chromatin-bound proteins isolated by the yChEFs procedure ^49, 50^ from the wild-type and *puf3Δ* strains grown in YPD medium at 30°C were analyzed by western blot using specific antibodies against Rpb4, H3 histone used as a positive control and Pgk1 as a negative control of cytoplasmic contamination. **B)** Whole-cell extract and chromatin-bound proteins isolated by the yChEFs procedure ^49^ from the Puf3-TAP tagged wild-type and *rpb4Δ* strains grown in YPD medium at 30°C were analyzed by western blot using specific antibodies against Rpb4, Rpb3 and TAP, and against H3 histone used as a positive control and Pgk1 employed as a negative control of cytoplasmic contamination.

Although Puf3 has been described as a cytosolic protein located on the mitochondrial surface ^40^, we also explored if Puf3 could be associated with chromatin. After following the yChEFs procedure ^49, 50^, Puf3-TAP was analysed by western blot in chromatin-enriched fractions of the *rpb4Δ* and wild-type strains expressing Puf3-TAP. Strikingly Puf3 was clearly detected in association with chromatin in a wild-type strain (Figure 7B). Notably, the association of Puf3 with chromatin seemed to depend on Rpb4 as chromatin-associated Puf3 decreased in an *rpb4Δ* strain, and the lower migrating wild-type Puf3 band mainly appeared. In addition, the faster migrating band also appeared but to a lesser extent.

In order to go further into the chromatin association of Puf3, we performed ChIP-Seq experiment of Puf3-TAP in both a wild-type and an *rpb4Δ* strain using a no-tag strain as the negative control. Unfortunately, the analysis of three biological replicates showed a similar IP signal than for the no-tag strain, which suggests that Puf3 did not bind likely chromatin tightly enough to be detected by chromatin immunoprecipitation (Figure S4).

Taken together, these results indicated that Puf3 integrity depends on Rpb4-bound to the chromatin-associated RNA pol II and likely, not on the Rpb4-mRNA association. Furthermore, our results also demonstrated that Puf3 associates to chromatin in a similar way.

### Rpb4 and Puf3 association with mRNAs depends on one another

As demonstrated above, Rpb4 and Puf3 genetically and physically interact. The Rpb4-RIP experiments showed that Rpb4 binds to Puf3 mRNAs, probably by modulating their stability. To decipher if Rpb4 also modulated the Puf3-mRNA interaction, we performed UV-crosslinking and protein-mRNA isolation experiments ^13^ to analyze the global association of Puf3 and Rpb4 to mRNAs by western blot in a wild-type strain and the *rpb4Δ* mutant strains expressing a TAP-tagged version of Puf3. As shown in Figure 8A, only a very weak Puf3-mRNA association seemed to occur in the cells lacking Rpb4. We also investigated whether lack of Puf3 could influence the Rpb4-mRNA association in a wild-type strain and *puf3Δ* mutant strains, and demonstrated that the association of Rpb4 with mRNA clearly diminished when Puf3 was lacking (Figure 8B).

**Figure 8:**
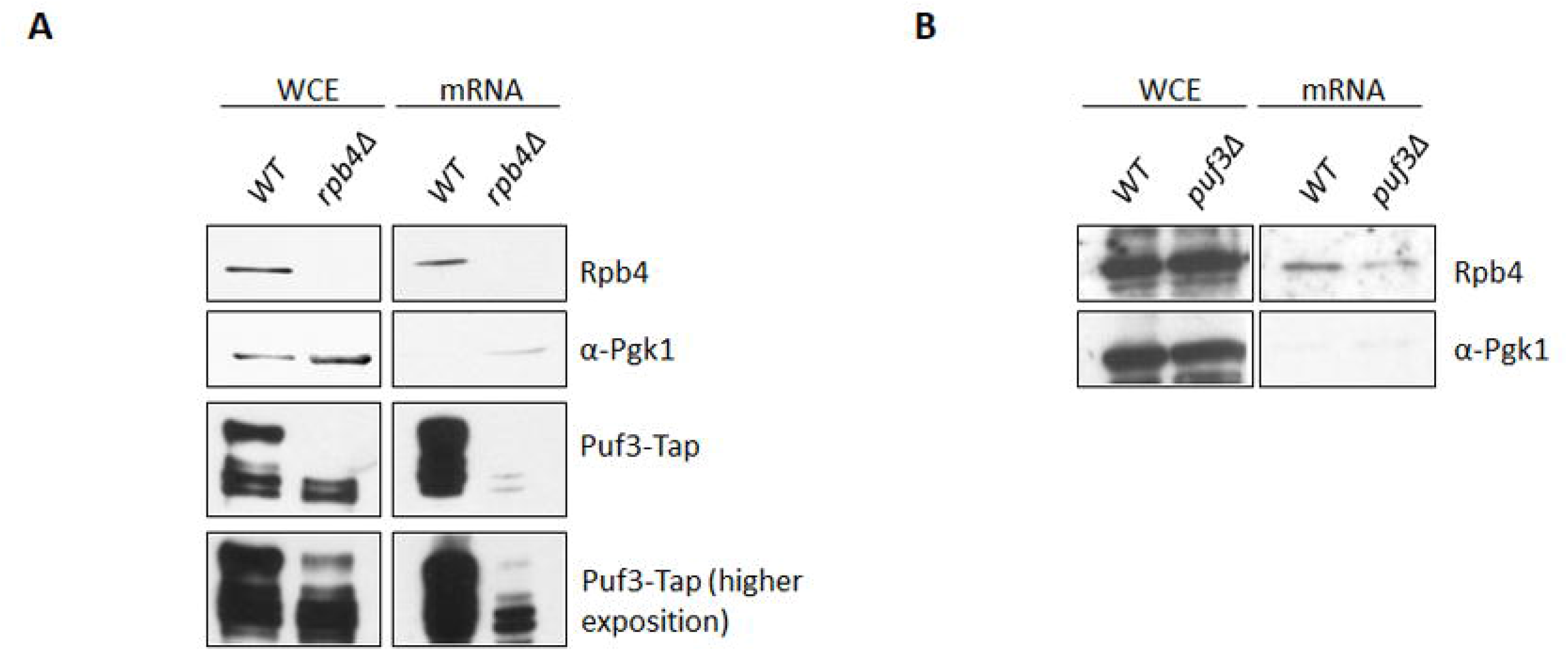
Rpb4 and Puf3 associations with mRNAs depend on each other. **A)** Western blot of both whole-cell extracts and oligo-dT purified mRNAs after exposure to 1200 mJ/cm^2^ of 254 nm UV light to crosslink proteins to mRNAs in the Puf3-TAP tagged wild-type and *rpb4Δ* strains grown in SD medium at 30°C. Western blots were carried out with specific antibodies against Rpb4, Tap-tag and Pgk1 used as a control of cytoplasmic protein (unbound to mRNA). **B)** Rpb4 association with mRNAs in a wild-type and *puf3Δ* strains analyzed under similar conditions to those indicated in **A**.

Those data indicate that the Rpb4 association with mRNA depends on Puf3, and the Puf3 association with mRNA depends on Rpb4, likely for a group of mRNAs.

## Discussion

mRNA synthesis and decay are interconnected and coregulated converting gene expression in a circular system, which changes the classic linear flow view of the Central Dogma of Molecular Biology ^8^. This mRNA crosstalk involves not only transcriptional machinery elements that impact mRNA decay, but also others from mRNA decay machinery that influence mRNA synthesis. Only Rpb4, a subunit of RNA pol II, has been clearly shown to be a transcriptional machinery element that participates in the entire mRNA life cycle ^5, 7, 12–14, 17^. This role by Rpb4 seems to be also participated by Rpb7, at least in part, as a dissociable dimer from the rest of the enzyme ^15, 18, 62^. We herein demonstrate by RIP-Seq experiments that Rpb4 specifically associates with a large mRNA population. A group of these mRNAs are also target of RBP Puf3 and correspond to nuclear genes for mitochondrial proteins. Our results demonstrate that Rpb4 and Puf3 physically, genetically and functionally interact, and suggest that Rpb4 and Puf3 cooperate to regulate their mRNA association and mRNA stability. Finally, our data indicate that Rpb4 could be a central transcriptional machinery component in the interplay between mRNA synthesis and degradation.

Rpb4-mRNA binding has been previously reported for some specific mRNAs ^7, 14, 17^ or more generally ^13^. Notably, our results demonstrate that Rpb4 associates with almost 1500 mRNAs, and point out that Rpb4 is a more general “imprinting element” than previously shown ^7, 14^, which falls in line with the proposed role of Rpb4 generally modulating mRNA imprinting and stability ^13^. Accordingly, the functional analysis of the GO categories identified different biological processes, the most significantly related ones to mitochondria, but also others that had significant p-values and corresponded to transmembrane transport or metabolic processes. However, the RIP experiments did not detect the specific class of mRNAs that have been previously proposed to be the targets of Rpb4, namely those for RP and RiBi genes ^14^. Although these data seem controversial, they could result from the fact that these targets have been established according to the role of Rpb4 in mRNA stability, and not directly based on its mRNA association ^7, 14^. However, we cannot rule out that Rpb4 imprints all or most mRNAs with different affinities, and the sensitivity of the applied technique was the main limitation to detect those weakly imprinted Rpb4-mRNA targets.

The most significantly identified GO categories establish a specific group of Rpb4-bound mRNA targets related to mitochondrion and mitochondrial translation, which correlate mainly with Pumilio family RPB Puf3-bound mRNAs ^37, 38, 40–42, 44, 63^. Accordingly, these Rpb4/Puf3 common targets contain the previously described 3’UTR motifs for Puf3-mRNA binding ^44^. In line with this, the physical and genetic interactions shown herein between Rpb4 and Puf3 corroborate these data. These results indicate that Rpb4 cooperates with RBPs, likely to modulate the mRNA turnover of a specific class of mRNAs, as our data also suggest by showing physical interactions between Rpb4 and Puf2 or Nsr1 ^35, 36, 56, 64, 65^. Puf2 is also a Pumilio family RBP protein that participates in destabilizing ribosome biogenesis, RiBi, mRNAs ^66^, whereas Nsr1 is a RiBi component ^65^ whose mRNA turnover depends on Rpb4 ^14^. Interestingly, these data point out the interplay between Rpb4-RPBs cooperation and the role of Rpb4 in mRNA decay of RiBi components, as previously demonstrated ^7, 14^.

A role for Rpb4 as a key element in mRNA synthesis and decay has been clearly established ^5, 7, 10, 12–14, 17^. Accordingly, the hypothesis that Rpb4 imprints mRNAs and decreases their stability ^13, 17^ falls in line with the fact that Rpb4-bound mRNAs decay faster. However, the change in mRNA stability by lack of Rpb4 does not differ between Rpb4-bound mRNAs, and other mRNAs indicating a general effect of Rpb4 on mRNA degradation and not to specific effects on given groups of mRNAs. Our results also suggest that the simultaneous binding of Rpb4 and Puf3 to mRNAs is not necessary for the effect of Rpb4 on Puf3 mRNA targets.

Our results demonstrate for the first time, that Puf3 binds chromatin, albeit weakly or transiently, and likely imprints mRNAs co-transcriptionally. In addition, both processes seem to depend on Rpb4. In fact lack of Rpb4 affected the amount of both chromatin-associated Puf3 and Puf3-mRNA binding. However, on the contrary, chromatin association of Rpb4 is independent of Puf3, since lack of Puf3 does not alter the amount of Rpb4 bound to chromatin.

Strikingly, Rpb4 is necessary to maintain Puf3 integrity as lack of Rpb4 affected Puf3 migration by leading to a faster-migrating band (~100 kDa) to accumulate (Figure 5). This proteolysis seems to correspond to the amino terminal domain of the protein as the anti-PAP antibody used to detect Puf3-TAP recognizes the C-terminal domain. The amino terminal domain has been shown to be phosphorylated under glucose deprivation by activating the translation of pre-accumulated mRNAs, and quickly favoring mitochondria biogenesis under respiration conditions ^38^. This function for the amino terminal domain of the protein accounts for Puf3 switching the fate of their mRNA targets from degradation to translation to facilitate cell acclimation to carbon-source availability and to fermentation/respiration conditions ^37, 38, 41^. Moreover, Puf3 integrity depends on Rpb4 associated with chromatin-bound RNA pol II, and not on the Rpb4-mRNA association, as the Puf3 faster-migrating band does not appear when cells express a Rpb2-Rpb4 fusion protein that remains associated mainly with chromatin ^17^. However, we were unable to rule out that the small fraction of Rpb2-Rpb4 or free Rpb4 that imprinted mRNAs in *rpb4Δ* cells expressing Rpb2-Rpb4 fusion protein could compensate, at least in part, the Puf3 integrity. It is worth noting that the faster-migrating Puf3 band bound chromatin less efficiently than the wild-type band (Figure 7), and did not seem to bind mRNAs efficiently (Figure 8). Accordingly, it is tempting to speculate that Rpb4 associates with Puf3 and stabilizes the protein to allow its efficient association to chromatin and its co-transcriptional mRNA imprinting. Furthermore, we also hypothesize that the low stability of Puf3 mRNA targets on the cells growing with glucose as a carbon source could be reached by the concerted action of Puf3 with Rpb4 because Rpb4-imprinted mRNAs decay faster and lack of Rpb4 increases mRNA stability.

Therefore, we speculated that Puf3-mRNA imprinting depended on Rpb4, probably as a consequence of Puf3 integrity. Similarly, Rpb4 mRNA imprinting would also be Puf3-dependent. We speculated that this dependence had to occur for commonly Rpb4-and Puf3-bound mRNAs, and could similarly take place for different classes of mRNAs with the concerted participation of Rpb4 and other RBPs based on the observed physical interaction between Rpb4 and other RPBs.

Taken together, our results showed that Rpb4 and Puf3 cooperated to regulate their mRNA association and mRNA degradation, and likely mRNA imprinting, at least for a common set of mRNAs associated with mitochondria. This could also occur for Rpb4 in concert with other RPBs.

Our results herein allowed us to hypothesize a functional model for Rpb4-Puf3 cooperation in mRNA stability. Puf3 bound to chromatin in an Rpb4-dependent manner, which also determined co-transcriptional Puf3-mRNA imprinting and influenced cytoplasmic mRNA degradation. Furthermore, Rpb4 imprinted a large set of mRNAs, part of which are common with Puf3. Regardless of this, Rpb4 was able to modulate the cytoplasmic functions of mRNA degradation machinery. In line with this, the cooperation between Rpb4 and mRNA decay machinery, such as Pat1-Lsm1/7, has been proposed to mediate mRNA decay ^14^. Rpb4-bound mRNAs have lower half-lives than overall mRNA and lack of Rpb4 increases mRNA stability. However, this increase did not seem to specifically occur for the set of Rpb4-bound mRNAs. This could be due to Rpb4’s very low level of imprinting in the rest of the transcriptome or because the effect of Rpb4 on global mRNA stability does not directly depend on mRNA imprinting. In any case, mRNA imprinting by Rpb4 seems related to mRNA stability if we consider that the mutants affecting the dissociation of Rpb4, or of dimer Rpb4/7 from RNA pol II, are also associated with lower mRNA imprinting and higher mRNA stability ^13, 17^. Notably however, Puf3-mRNA imprinting impacts mRNA stability as the stability of all Puf3-bound mRNAs increases upon Puf3 deletion ^38^, and also because the increase in mRNA stability provoked by lack of Rpb4 is attenuated for the Puf3-bound mRNAs. We speculated that the alteration of Puf3 integrity provoked by lack of Rpb4, which diminished Puf3-mRNA imprinting, negatively affected the increased Puf3-bound mRNA stability to a lower extent than for either other Rpb4-bound mRNAs or overall mRNAs, likely by the abnormal concerted action of Puf3 and mRNA degradation machinery.

Furthermore, by considering that the concerted action among Puf3, Pab1 and the Ccr4-Not complex has been shown to act on mRNA poly(A) tails to regulate eukaryotic mRNA stability and translation ^11, 67, 68^, we also speculated that Rpb4-Puf3 cooperation could serve to influence Ccr4-Not activity in mRNA degradation. Notably, the Puf3 binding motif is overrepresented in the most highly enriched Ccr4 targets ^69^. Similarly to that occurs for Rpb4 over Puf3 integrity, Rpb4 could act on other elements of the mRNA degradation machinery and acting in concert on mRNA stability.

Finally, it would be interesting to analyze the possible mRNA targets of Rpb4 under different growth conditions to establish the most precise role of this protein as a central transcriptional machinery element that connects the mRNA degradation process.

## ACKNOWLEDGMENTS

We thank the “Servicios Centrales de Apoyo a la Investigación (SCAI)” of the University of Jaen for technical support. We thank Olga Calvo for sharing with us the *S. pombe* spike in.

## REFERENCES

1. Pérez-Ortín JE, Alepuz P, Chávez S, Choder M. Eukaryotic mRNA decay: methodologies, pathways, and links to other stages of gene expression. J Mol Biol 2013; 425:3750–75.

2. Sun M, Schwalb B, Schulz D, Pirkl N, Etzold S, Lariviere L, et al. Comparative dynamic transcriptome analysis (cDTA) reveals mutual feedback between mRNA synthesis and degradation. Genome Res 2012; 22:1350–9.

3. Braun KA, Young ET. Coupling mRNA synthesis and decay. Mol Cell Biol 2014; 34:4078–87.

4. Begley V, Corzo D, Jordán-Pla A, Cuevas-Bermúdez A, Miguel-Jiménez L, Pérez-Aguado D, et al. The mRNA degradation factor Xrn1 regulates transcription elongation in parallel to Ccr4. Nucleic Acids Res 2019; 47:9524–41.

5. Choder M. mRNA imprinting: Additional level in the regulation of gene expression. Cell Logist 2011; 1:37–40.

6. Dahan N, Choder M. The eukaryotic transcriptional machinery regulates mRNA translation and decay in the cytoplasm. Biochim Biophys Acta 2013; 1829:169–73.

7. Goler-Baron V, Selitrennik M, Barkai O, Haimovich G, Lotan R, Choder M. Transcription in the nucleus and mRNA decay in the cytoplasm are coupled processes. Genes Dev 2008; 22:2022–7.

8. Haimovich G, Medina DA, Causse SZ, Garber M, Millán-Zambrano G, Barkai O, et al. Gene expression is circular: factors for mRNA degradation also foster mRNA synthesis. Cell 2013; 153:1000–11.

9. Harel-Sharvit L, Eldad N, Haimovich G, Barkai O, Duek L, Choder M. RNA polymerase II subunits link transcription and mRNA decay to translation. Cell 2010; 143:552–63.

10. Shalem O, Groisman B, Choder M, Dahan O, Pilpel Y. Transcriptome kinetics is governed by a genome-wide coupling of mRNA production and degradation: a role for RNA Pol II. PLoS Genet 2011; 7:e1002273.

11. Moqtaderi Z, Geisberg JV, Struhl K. Extensive structural differences of closely related 3’ mRNA isoforms: links to Pab1 binding and mRNA stability. Mol Cell 2018; 72:849–61 e6.

12. Farago M, Nahari T, Hammel C, Cole CN, Choder M. Rpb4p, a subunit of RNA polymerase II, mediates mRNA export during stress. Mol Biol Cell 2003; 14:2744–55.

13. Garrido-Godino AI, García-López MC, García-Martínez J, Pelechano V, Medina DA, Pérez-Ortín JE, et al. Rpb1 foot mutations demonstrate a major role of Rpb4 in mRNA stability during stress situations in yeast. Biochim Biophys Acta 2016; 1859:731–43.

14. Lotan R, Bar-On VG, Harel-Sharvit L, Duek L, Melamed D, Choder M. The RNA polymerase II subunit Rpb4p mediates decay of a specific class of mRNAs. Genes Dev 2005; 19:3004–16.

15. Lotan R, Goler-Baron V, Duek L, Haimovich G, Choder M. The Rpb7p subunit of yeast RNA polymerase II plays roles in the two major cytoplasmic mRNA decay mechanisms. J Cell Biol 2007; 178:1133–43.

16. Kumar D, Varshney S, Sengupta S, Sharma N. A comparative study of the proteome regulated by the Rpb4 and Rpb7 subunits of RNA polymerase II in fission yeast. J Proteomics 2019; 199:77–88.

17. Duek L, Barkai O, Elran R, Adawi I, Choder M. Dissociation of Rpb4 from RNA polymerase II is important for yeast functionality. PLoS One 2018; 13:e0206161.

18. Selitrennik M, Duek L, Lotan R, Choder M. Nucleocytoplasmic shuttling of the Rpb4p and Rpb7p subunits of *Saccharomyces cerevisiae* RNA polymerase II by two pathways. Eukaryot Cell 2006; 5:2092–103.

19. Edwards AM, Kane CM, Young RA, Kornberg RD. Two dissociable subunits of yeast RNA polymerase II stimulate the initiation of transcription at a promoter in vitro. J Biol Chem 1991; 266:71–5.

20. Sampath V, Balakrishnan B, Verma-Gaur J, Onesti S, Sadhale PP. Unstructured N terminus of the RNA polymerase II subunit Rpb4 contributes to the interaction of Rpb4.Rpb7 subcomplex with the core RNA polymerase II of *Saccharomyces cerevisiae*. J Biol Chem 2008; 283:3923–31.

21. Schulz D, Pirkl N, Lehmann E, Cramer P. Rpb4 subunit functions mainly in mRNA synthesis by RNA polymerase II. J Biol Chem 2014; 289:17446–52.

22. Allepuz-Fuster P, O’Brien MJ, González-Polo N, Pereira B, Dhoondia Z, Ansari A, et al. RNA polymerase II plays an active role in the formation of gene loops through the Rpb4 subunit. Nucleic Acids Res 2019; 47:8975–898.

23. Forget A, Chartrand P. Cotranscriptional assembly of mRNP complexes that determine the cytoplasmic fate of mRNA. Transcription 2011; 2:86–90.

24. Villanyi Z, Ribaud V, Kassem S, Panasenko OO, Pahi Z, Gupta I, et al. The Not5 subunit of the Ccr4-Not complex connects transcription and translation. PLoS Genet 2014; 10:e1004569.

25. Li T, De Clercq N, Medina DA, Garre E, Sunnerhagen P, Perez-Ortin JE, et al. The mRNA cap-binding protein Cbc1 is required for high and timely expression of genes by promoting the accumulation of gene-specific activators at promoters. Biochim Biophys Acta 2016; 1859:405–19.

26. Glisovic T, Bachorik JL, Yong J, Dreyfuss G. RNA-binding proteins and post-transcriptional gene regulation. FEBS letters 2008; 582:1977–86.

27. Keene JD. RNA regulons: coordination of post-transcriptional events. Nature Reviews Genetics 2007; 8:533.

28. Halbeisen RE, Galgano A, Scherrer T, Gerber AP. Post-transcriptional gene regulation: from genome-wide studies to principles. Cellular and molecular life sciences 2008; 65:798.

29. Jiang H, Xu L, Wang Z, Keene J, Gu Z. Coordinating expression of RNA binding proteins with their mRNA targets. Scientific reports 2014; 4:7175.

30. Baltz AG, Munschauer M, Schwanhäusser B, Vasile A, Murakawa Y, Schueler M, et al. The mRNA-bound proteome and its global occupancy profile on protein-coding transcripts. Molecular cell 2012; 46:674–90.

31. Mitchell SF, Jain S, She M, Parker R. Global analysis of yeast mRNPs. Nat Struct Mol Biol 2013; 20:127–33.

32. Khong A, Parker R. The landscape of eukaryotic mRNPs. RNA 2020; 26:229–39.

33. Holmqvist E, Vogel J. RNA-binding proteins in bacteria. Nat Rev Microbiol 2018; 16:601–15.

34. Klass DM, Scheibe M, Butter F, Hogan GJ, Mann M, Brown PO. Quantitative proteomic analysis reveals concurrent RNA-protein interactions and identifies new RNA-binding proteins in *Saccharomyces cerevisiae*. Genome Res 2013; 23:1028–38.

35. Quenault T, Lithgow T, Traven A. PUF proteins: repression, activation and mRNA localization. Trends in cell biology 2011; 21:104–12.

36. Wickens M, Bernstein DS, Kimble J, Parker R. A PUF family portrait: 3’ UTR regulation as a way of life. TRENDS in Genetics 2002; 18:150–7.

37. Lee CD, Tu BP. Glucose-Regulated phosphorylation of the PUF protein Puf3 regulates the translational fate of its bound mRNAs and association with RNA granules. Cell Rep 2015; 11:1638–50.

38. Miller MA, Russo J, Fischer AD, Lopez Leban FA, Olivas WM. Carbon source-dependent alteration of Puf3p activity mediates rapid changes in the stabilities of mRNAs involved in mitochondrial function. Nucleic Acids Res 2014; 42:3954–70.

39. Wang X, Voronina E. Diverse roles of PUF proteins in germline stem and progenitor cell development in *C. elegans*. Front Cell Dev Biol 2020; 8:29.

40. García-Rodríguez LJ, Gay AC, Pon LA. Puf3p, a Pumilio family RNA binding protein, localizes to mitochondria and regulates mitochondrial biogenesis and motility in budding yeast. J Cell Biol 2007; 176:197–207.

41. Lapointe CP, Stefely JA, Jochem A, Hutchins PD, Wilson GM, Kwiecien NW, et al. Multi-omics reveal specific targets of the RNA-binding protein Puf3p and its orchestration of mitochondrial biogenesis. Cell Syst 2018; 6:125–35 e6.

42. Saint-Georges Y, Garcia M, Delaveau T, Jourdren L, Le Crom S, Lemoine S, et al. Yeast mitochondrial biogenesis: a role for the PUF RNA-binding protein Puf3p in mRNA localization. PloS one 2008; 3:e2293.

43. García-López MC, Mirón-García MC, Garrido-Godino AI, Mingorance C, Navarro F. Overexpression of *SNG1* causes 6-azauracil resistance in *Saccharomyces cerevisiae*. Curr Genet 2010; 56:251–63.

44. Gupta I, Clauder-Munster S, Klaus B, Jarvelin AI, Aiyar RS, Benes V, et al. Alternative polyadenylation diversifies post-transcriptional regulation by selective RNA-protein interactions. Mol Syst Biol 2014; 10:719.

45. Wilkening S, Pelechano V, Jarvelin AI, Tekkedil MM, Anders S, Benes V, et al. An efficient method for genome-wide polyadenylation site mapping and RNA quantification. Nucleic Acids Res 2013; 41:e65.

46. Anders S, Huber W. Differential expression analysis for sequence count data. Genome Biol 2010; 11:R106.

47. Mirón-García MC, Garrido-Godino AI, García-Molinero V, Hernández-Torres F, Rodríguez-Navarro S, Navarro F. The prefoldin Bud27 mediates the assembly of the eukaryotic RNA polymerases in an Rpb5-dependent manner. PLoS Genet 2013; 9:e1003297.

48. Garrido-Godino AI, García-López MC, Navarro F. Correct assembly of RNA polymerase II Depends on the foot domain and Is required for multiple steps of transcription in *Saccharomyces cerevisiae*. Mol Cell Biol 2013; 33:3611–26.

49. Cuevas-Bermúdez A, Garrido-Godino A, Navarro F. A novel yeast chromatin-enriched fractions purification approach, yChEFs, for the chromatin-associated protein analysis used for chromatin-associated and RNA-dependent chromatin-associated proteome studies from *Saccharomyces cerevisiae*. Gene Reports 2019; 16:100450.

50. Cuevas-Bermúdez A, Garrido-Godino AI, Gutiérrez-Santiago F, Navarro F. A Yeast Chromatin-enriched Fractions Purification Approach, yChEFs, from *Saccharomyces cerevisiae*. Bio-protocol 2020; 10:e3471.

51. Coulombe C, Poitras C, Nordell-Markovits A, Brunelle M, Lavoie MA, Robert F, et al. VAP: a versatile aggregate profiler for efficient genome-wide data representation and discovery. Nucleic Acids Res 2014; 42:W485–93.

52. Love MI, Huber W, Anders S. Moderated estimation of fold change and dispersion for RNA-seq data with DESeq2. Genome Biol 2014; 15:550.

53. Szklarczyk D, Morris JH, Cook H, Kuhn M, Wyder S, Simonovic M, et al. The STRING database in 2017: quality-controlled protein-protein association networks, made broadly accessible. Nucleic Acids Res 2017; 45:D362–D8.

54. Kershaw CJ, Costello JL, Talavera D, Rowe W, Castelli LM, Sims PF, et al. Integrated multi-omics analyses reveal the pleiotropic nature of the control of gene expression by Puf3p. Scientific reports 2015; 5:15518.

55. Wilinski D, Buter N, Klocko AD, Lapointe CP, Selker EU, Gasch AP, et al. Recurrent rewiring and emergence of RNA regulatory networks. Proceedings of the National Academy of Sciences 2017; 114:E2816–E25.

56. Gerber AP, Herschlag D, Brown PO. Extensive association of functionally and cytotopically related mRNAs with Puf family RNA-binding proteins in yeast. PLoS biology 2004; 2:e79.

57. Freeberg MA, Han T, Moresco JJ, Kong A, Yang YC, Lu ZJ, et al. Pervasive and dynamic protein binding sites of the mRNA transcriptome in *Saccharomyces cerevisiae*. Genome Biol 2013; 14:R13.

58. Hogan DJ, Riordan DP, Gerber AP, Herschlag D, Brown PO. Diverse RNA-binding proteins interact with functionally related sets of RNAs, suggesting an extensive regulatory system. PLoS Biol 2008; 6:e255.

59. Garcia-Martinez J, Aranda A, Perez-Ortin JE. Genomic run-on evaluates transcription rates for all yeast genes and identifies gene regulatory mechanisms. Mol Cell 2004; 15:303–13.

60. Escobar-Henriques M, Daignan-Fornier B. Transcriptional regulation of the yeast gmp synthesis pathway by its end products. Journal of Biological Chemistry 2001; 276:1523–30.

61. Wei W, Hennig BP, Wang J, Zhang Y, Piazza I, Pareja Sanchez Y, et al. Chromatin-sensitive cryptic promoters putatively drive expression of alternative protein isoforms in yeast. Genome Res 2019; 29:1974–84.

62. Choder M. Rpb4 and Rpb7: subunits of RNA polymerase II and beyond. Trends Biochem Sci 2004; 29:674–81.

63. Gadir N, Haim-Vilmovsky L, Kraut-Cohen J, Gerst JE. Localization of mRNAs coding for mitochondrial proteins in the yeast *Saccharomyces cerevisiae*. RNA 2011; 17:1551–65.

64. Hogan GJ, Brown PO, Herschlag D. Evolutionary conservation and diversification of Puf RNA binding proteins and their mRNA targets. PLoS biology 2015; 13:e1002307.

65. Lee WC, Zabetakis D, Melese T. NSR1 is required for pre-rRNA processing and for the proper maintenance of steady-state levels of ribosomal subunits. Mol Cell Biol 1992; 12:3865–71.

66. Fischer AD, Olivas WM. Multiple Puf proteins regulate the stability of ribosome biogenesis transcripts. RNA Biol 2018; 15:1228–43.

67. Webster MW, Stowell JA, Passmore LA. RNA-binding proteins distinguish between similar sequence motifs to promote targeted deadenylation by Ccr4-Not. eLife 2019; 8.

68. Olivas W, Parker R. The Puf3 protein is a transcript-specific regulator of mRNA degradation in yeast. EMBO J 2000; 19:6602–11.

69. Miller JE, Zhang L, Jiang H, Li Y, Pugh BF, Reese JC. Genome-Wide Mapping of Decay Factor-mRNA Interactions in Yeast Identifies Nutrient-Responsive Transcripts as Targets of the Deadenylase Ccr4. G3 (Bethesda) 2018; 8:315–30.

70. Riordan DP, Herschlag D, Brown PO. Identification of RNA recognition elements in the Saccharomyces cerevisiae transcriptome. Nucleic Acids Res 2011; 39:1501–9.

