## Supplementary Material for "Rpb4 and Puf3 imprint and post-transcriptionally control the stability of a common set of mRNAs in yeast"

**Table S1.** ***S. cerevisiae*** **strains**

| Strain | Genotype | Origin |
|---|---|---|
| BY4741 | *MATa his3-Δ1 leu2-Δ0 met15-Δ0 ura3-Δ0* | Euroscarf |
| Y223 | *MATa pep4::HIS3/prb1::LEU2 prc1::HISG can1 ade2 trp1 ura3 his3 leu2-3,112 RPB4::TAP::TRP1* | M. Choder |
| YFN116 | *MATα his3-Δ200 leu2-3,112 trp1-Δ63 ura3-52 rpb1-Δ187::HIS3 + pYEB220 (2μm LEU2 RPB1)* | [1] |
| YFN104 | *MATα his3-Δ200 leu2-3,112 trp1-Δ63 ura3-52 rpb1-Δ187::HIS3 + pYEB220-84 (2μm LEU2 rpb1-84)* | [1] |
| YFN518 | *MATα his3-Δ200 leu2-3,112 trp1-Δ63 ura3-52 rpb1-Δ187::HIS3 + pYEB220 (2μm LEU2 RPB1) RPB4::TAP::TRP1* | This work |
| YFN519 | *MATα his3-Δ200 leu2-3,112 trp1-Δ63 ura3-52 rpb1-Δ187::HIS3 + pYEB220-84 (2μm LEU2 rpb1-84) RPB4::TAP::TRP1* | This work |
| YFN665 | *MATα ade2 leu2-3,112 ura3-52 trp1* rpb1-Δ187::kanMX4 + pYEB220 (2mm LEU) puf3::TAP::URA* | This work |
| YFN667 | *MATα ade2 leu2-3,112 ura3-52 trp1* rpb1-Δ187::kanMX4 + pYEB220-84 (2μm LEU2 rpb1-84) puf3::TAP::URA* | This work |
| YFN678 | *MATa ade2 arg4 leu2-3,112 trp1-289 ura3-52 puf3::TAP::URA rpb4::kanMX4* | This work |
| YFN749 | *MATα his3Δ1 leu2Δ0 lys2Δ0 ura3Δ0 rpb4Δ::HIS3 puf3Δ::kanMX4* | This work |
| yMC798 | *MATa his3Δ1 leu2Δ0 met15Δ0 ura3Δ0 rpb2 ::KanMX6; pRS315-RPB2 (CEN, LEU2)* | [2] |
| yMC871 | *MATa his3Δ1 leu2Δ0 met15Δ0 ura3Δ0 rpb2 ::KanMX6; pRS315-RPB2-RPB4 (CEN, LEU2)* | [3] |
| YFN798 | *MATa his3Δ1 leu2Δ0 met15Δ0 ura3Δ0 rpb2 ::KanMX6; pRS315-RPB2 (CEN, LEU2) puf3::TAP::URA* | This work |
| YFN799 | *MATa his3Δ1 leu2Δ0 met15Δ0 ura3Δ0 rpb2 ::KanMX6; pRS315-* | This work |

| | | |
|---|---|---|
| | *RPB2-RPB4 (CEN, LEU2) puf3::TAP::URA* | |
| YFN800 | *MATa his3Δ1 leu2Δ0 met15Δ0 ura3Δ0 rpb2 ::KanMX6; pRS315-RPB2 (CEN, LEU2) puf3::TAP::URA rpb4::HIS3* | This work |
| Not1-TAP | *MATa his3Δ1 leu2Δ0 met15Δ0 ura3Δ0 not1::TAP::HIS3* | [4] |
| Nrd1-TAP | *MATa ade2 arg4 leu2-3,112 trp1-289 ura3-52 nrd1::TAP::URA3* | [5] |
| Nsr1-TAP | *MATa ade2 arg4 leu2-3,112 trp1-289 ura3-52 nsr1::TAP::URA3* | [5] |
| Imd2-TAP | *MATa His3Δ1 Leu2Δ0 met15Δ0 ura3Δ0 imd2::TAP::HIS3MX6* | Open Biosystems |
| Rpb3-TAP | *MATa his3Δ1 leu2Δ0 met15Δ0 ura3Δ0 rpb3::TAP::HIS3Mx6* | Open Biosystems |
| Pub1-TAP | *MATa ade2 arg4 leu2-3,112 trp1-289 ura3-52 pub1::TAP::URA3* | [5] |
| Puf2-TAP | *MATa ade2 arg4 leu2-3,112 trp1-289 ura3-52 puf2::TAP::URA3* | [5] |
| Puf3-TAP | *MATa ade2 arg4 leu2-3,112 trp1-289 ura3-52 puf3::TAP::URA* | [5] |
| Puf4-TAP | *MATa ade2 arg4 leu2-3,112 trp1-289 ura3-52 puf4::TAP::URA3* | [5] |
| Puf5-TAP | *MATa ade2 arg4 leu2-3,112 trp1-289 ura3-52 puf5::TAP::URA3* | [5] |
| Vts1-TAP | *MATa ade2 arg4 leu2-3,112 trp1-289 ura3-52 vts1::TAP::URA3* | [5] |

**Table S2. Oligonucleotides used**

| | | |
|---|---|---|
| *18S*-501 | CATGGCCGTTCTTAGTTGGT | ORF |
| *18S*-301 | ATTGCCTCAAACTTCCATCG | ORF |
| *LYS20*-501 | ATCACCGGGTTCTGTGCATT | ORF |
| *LYS20*-301 | GCGTTCCAGCCAGTTAGTCT | ORF |
| *HHF1-501* | GCCAAGCGTCACAGAAAG AT | ORF |
| *HHF1-301* | GATGACGGATTCCAAGAAGG | ORF |
| *MRP1-501* | CTGAGATGACTGCCAAGCAA | ORF |
| *MRP1-301* | TTCTCGTTTCCCAAATACGC | ORF |
| *DRF1-501* | GGGGTGGCGAAGTTTATAGTCA | ORF |
| *DRF1-301* | TCACAGTCTGTTTCGGGCAA | ORF |
| *OAC1-501* | CAGCAAAGAACATTTTGGTCA | ORF |
| *OAC1-301* | ATCCCATGGGTTCATAACGA | ORF |
| *URA1-501* | GCCAATGTTCGTGCATTTTA | ORF |
| *URA1-301* | TCTTGACGCCCTCTTTTTGT | ORF |
| *RPL3-501* | AAGAAGGCTCATTTGGCTG | ORF |
| *RPL3-301* | AACACCTTCGAAACCGTGAC | ORF |
| *PYK1-4 (forw)* | CTATGGCTGAAACCGCTGTCATTG | ORF |
| *PYK1-4 (rev)* | CAGCTCTTGGGCATCTGGTAAC | ORF |
| *SCR1-501* | CGGCCGGGATAGCACATAT | ORF |
| *SCR1-301* | GACACACTCCATCCCCGAG | ORF |
| *PUF3-502* | TCTTCTTTAAGCGCCCTGTC | ORF, 5' region of the ORF |
| *PUF3-304* | TACTTCATTTGCCGTGGTGA | ORF, 5' region of the ORF |
| *PUF3-501* | TGCAACAAGATCAGTTCACCA | ORF, Internal region of the ORF |
| *PUF3-301* | CGACAACATTATTGGCCACAGT | ORF, Internal region of the ORF |
| *PUF3-503* | TGCTTTGAATCTGGAAGACG | ORF, 3' region of the ORF |
| *PUF3-305* | CCCTCACCCTCAGAGACATT | ORF, 3' region of the ORF |
| *PUF3-302* | AGCGTAAGAGCGCAAAGAT | 3' external region |
| *PUF3-303* | ACGGTTAGAAACAGGCTTGA | 3' external region |
| *RPB4-502* | TGCTTGCAATGGTTCAGAAG | 5' external region |
| *RPB4-301* | CCATTTTTGGTCGAATTTTG | 3' external region |

The primers used were as follows (all 5'-3')

**Table S3:** Functional categories (Biological Process). mRNAs identified by RIP-Seq of Rpb4 from *S. cerevisiae*. The analysis was performed using the STRING software [6] and the PPI enrichment p-value:<1.0e-16 was selected.

| Functional Category | FDR | Functional Category | FDR |
|---|---|---|---|
| Mitochondrial translation | 3.37e-07 | Cellular protein modification process | 0.0245 |
| Mitochondrial gene expression | 3.37e-07 | Response to chemical | 0.0258 |
| Transmembrane transport | 2.92e-06 | Localization | 0.0270 |
| Cellular process | 3.24e-06 | Anion transport | 0.0293 |
| Metal ion transport | 0.0013 | Organic anion transport | 0.0293 |
| Organonitrogen compound biosynthetic process | 0.0013 | Filamentous growth | 0.0293 |
| Metabolic process | 0.0014 | Cation transport | 0.0328 |
| Cellular lipid metabolic process | 0.0016 | Protein mannosylation | 0.0359 |
| Divalent inorganic cation transport | 0.0016 | Inorganic ion homeostasis | 0.0359 |
| Organonitrogen compound metabolic process | 0.0016 | Mitochondrial transport | 0.0362 |
| Ion transport | 0.0017 | Amino acid transmembrane transport | 0.0408 |
| Divalent metal ion transport | 0.0017 | Membrane lipid metabolic process | 0.0408 |
| Cell wall organization or biogenesis | 0.0017 | Lipid biosynthetic process | 0.0408 |
| Cell wall organization | 0.0018 | Cation homeostasis | 0.0408 |
| Transition metal ion transport | 0.0020 | Response to drug | 0.0449 |
| Lipid metabolic process | 0.0020 | Amino acid transport | 0.0461 |
| Transition metal ion homeostasis | 0.0021 | | |
| Organic substance metabolic process | 0.0022 | | |
| Metal ion homeostasis | 0.0024 | | |
| Drug transmembrane transport | 0.0036 | | |
| Cellular metal ion homeostasis | 0.0044 | | |
| Cellular protein metabolic process | 0.0044 | | |
| Cellular transition metal ion homeostasis | 0.0044 | | |
| Ion transmembrane transport | 0.0048 | | |
| Anion transmembrane transport | 0.0048 | | |
| Transport | 0.0060 | | |
| Cellular metabolic process | 0.0060 | | |
| Biosynthetic process | 0.0074 | | |
| Drug transport | 0.0085 | | |
| Primary metabolic process | 0.0085 | | |
| Cellular biosynthetic process | 0.0085 | | |
| Organic substance biosynthetic process | 0.0105 | | |
| Glycosylation | 0.0118 | | |
| Establishment of localization | 0.0135 | | |
| Mannosylation | 0.0152 | | |
| Protein glycosylation | 0.0180 | | |
| Growth | 0.0193 | | |
| Fungal-type cell wall organization or biogenesis | 0.0193 | | |
| Glycoprotein metabolic process | 0.0213 | | |
| Protein metabolic process | 0.0229 | | |
| Fungal-type cell wall organization | 0.0243 | | |

**Table S4:** Puf3-bound mRNAs. mRNAs were consider as Puf3 interactors (223) when they were detected in, at least, three of the five indicated datasets corresponding to Puf3-bound mRNAs [7-11].

| | | | | | |
|---|---|---|---|---|---|
| YGL043W | YGR112W | YHR147C | YOR150W | YKR085C | YGR165W |
| YKL053C-A | YPL059W | YNL131W | YGL107C | YPL097W | YER141W |
| YOR187W | YDR270W | YDR322W | YDR115W | YDR347W | YNL315C |
| YDR229W | YPL029W | YKR006C | YGR076C | YBR120C | YDR116C |
| YBR292C | YPL172C | YKL167C | YCR071C | YDR494W | YER093C-A |
| YJL071W | YMR098C | YJL205C | YKL155C | YNR020C | YNR045W |
| YLL013C | YLR382C | YLR277C | YKL170W | YNL137C | YJR122W |
| YGR161W-C | YPL040C | YJL102W | YMR225C | YBL107C | YNL098C |
| YNL145W | YHR005C-A | YNR022C | YBR185C | YLR218C | YOR205C |
| YBL059C-A | YHR011W | YNL026W | YPL132W | YDR237W | YJR113C |
| YJL209W | YGL129C | YPR067W | YBL038W | YMR089C | YLR357W |
| YOR267C | YMR166C | YJL096W | YOR354C | YDR194C | YNL306W |
| YJL133W | YHR159W | YPR166C | YBR192W | YBL080C | YGL143C |
| YMR054W | YGR238C | YDR511W | YDR375C | YNL252C | YLR067C |
| YGL064C | YJR135W-A | YDR041W | YNL122C | YLR439W | YDR430C |
| YDR028C | YLR202C | YKL003C | YCR046C | YPL260W | YBL008W |
| YCL051W | YER026C | YKL138C | YNL177C | YBR146W | YDR493W |
| YER122C | YOR020C | YMR158W | YJL062W-A | YMR193W | YNR037C |
| YNL030W | YNL070W | YDR286C | YJL046W | YBR262C | YKL087C |
| YNL197C | YBR268W | YNR017W | YBR122C | YER154W | YHR038W |
| YHR116W | YDR296W | YPL013C | YLL009C | YNR036C | YIL093C |
| YPR047W | YPL252C | YOR158W | YML025C | YFL036W | YBR172C |
| YBR179C | YML110C | YLR177W | YLR204W | YIL070C | YGR021W |
| YMR291W | YPR133W-A | YNL081C | YOL071W | YGR257C | YKL194C |
| YDR316W | YMR012W | YPR004C | YJL208C | YMR188C | YDR258C |
| YMR257C | YML009C | YGL068W | YEL020W-A | YDL182W | YEL050C |
| YGL236C | YLR069C | YHR119W | YHL004W | YMR157C | YOL033W |
| YDR337W | YLR390W | YDR451C | YHL014C | YMR287C | YOL023W |
| YHR059W | YJL104W | YDR127W | YJL063C | YNL083W | YHR091C |
| YAL039C | YML129C | YLR008C | YGR147C | YDL202W | YGL071W |
| YER058W | YOR286W | YMR286W | YLR259C | YNL073W | YIR021W |
| YCL036W | YJL143W | YGR028W | YGL226W | YCR003W | YDL044C |
| YGR062C | YDL045W-A | YJL054W | YPL173W | YIL098C | YLR090W |
| YHR090C | YPL183W-A | YPL072W | YBR251W | YGR084C | |
| YGR212W | YGR220C | YBR024W | YNL005C | YHR024C | |
| YDR079W | YNL185C | YDL107W | YBR258C | YHR169W | |
| YCR024C | YBR282W | YER050C | YNR040W | YDR405W | |
| YHR168W | YER182W | YOR201C | YNL284C | YMR024W | |

**Table S5:** Functional categories (Biological Process). Common interactors of Rpb4 (this study) and Puf3. mRNAs were consider as Puf3 interactors (223) when they were detected in three datasets of Puf3 bound mRNAs at least [7-11]. The analysis was performed using the STRING software [6] and the PPI enrichment p-value:<1.0e-16 was selected.

| Functional category | FDR | Functional category | FDR |
|---|---|---|---|
| Mitochondrial gene expression | 1.34e-59 | Trna aminoacylation for protein translation | 8.00e-06 |
| Mitochondrial translation | 1.82e-56 | Protein insertion into mitochondrial inner membrane | 8.00e-06 |
| Translation | 5.05e-34 | Nitrogen compound metabolic process | 1.60e-05 |
| Amide biosynthetic process | 3.80e-32 | Cellular process | 7.11e-05 |
| Peptide metabolic process | 4.09e-32 | Mitochondrial genome maintenance | 0.00011 |
| Organonitrogen compound biosynthetic process | 1.43e-20 | Cellular metabolic process | 0.00013 |
| Gene expression | 1.28e-15 | Metabolic process | 0.00032 |
| Cellular protein metabolic process | 1.45e-14 | Protein insertion into mitochondrial inner membrane fr | 0.00055 |
| Cellular nitrogen compound biosynthetic process | 3.44e-13 | Organic substance metabolic process | 0.00075 |
| Protein metabolic process | 5.43e-13 | Primary metabolic process | 0.0013 |
| Mitochondrial RNA metabolic process | 4.01e-12 | Protein transmembrane import into intracellular organe | 0.0014 |
| Cellular macromolecule biosynthetic process | 1.84e-11 | Cytochrome complex assembly | 0.0021 |
| Organonitrogen compound metabolic process | 3.62e-09 | Protein transmembrane transport | 0.0021 |
| Mitochondrion organization | 4.88e-09 | Mitochondrial RNA catabolic process | 0.0024 |
| Cellular nitrogen compound metabolic process | 1.94e-08 | Heme a biosynthetic process | 0.0024 |
| Cellular biosynthetic process | 2.01e-08 | Heme a metabolic process | 0.0024 |
| Mitochondrial transport | 2.66e-08 | Group I intron splicing | 0.0035 |
| Organic substance biosynthetic process | 3.29e-08 | Intracellular protein transmembrane transport | 0.0066 |
| Cellular macromolecule metabolic process | 4.72e-08 | Regulation of mitochondrial translation | 0.0083 |
| Trna aminoacylation for mitochondrial protein transla | 1.86e-07 | Mitochondrial respiratory chain complex assembly | 0.0239 |
| Mitochondrial transmembrane transport | 5.76e-07 | Protein import into mitochondrial matrix | 0.0254 |
| Macromolecule metabolic process | 1.19e-06 | Mitochondrial proton-transporting ATP synthase compl | 0.0320 |
| Protein targeting to mitochondrion | 1.55e-06 | Cytochrome c-heme linkage | 0.0484 |

**Heatpairs**

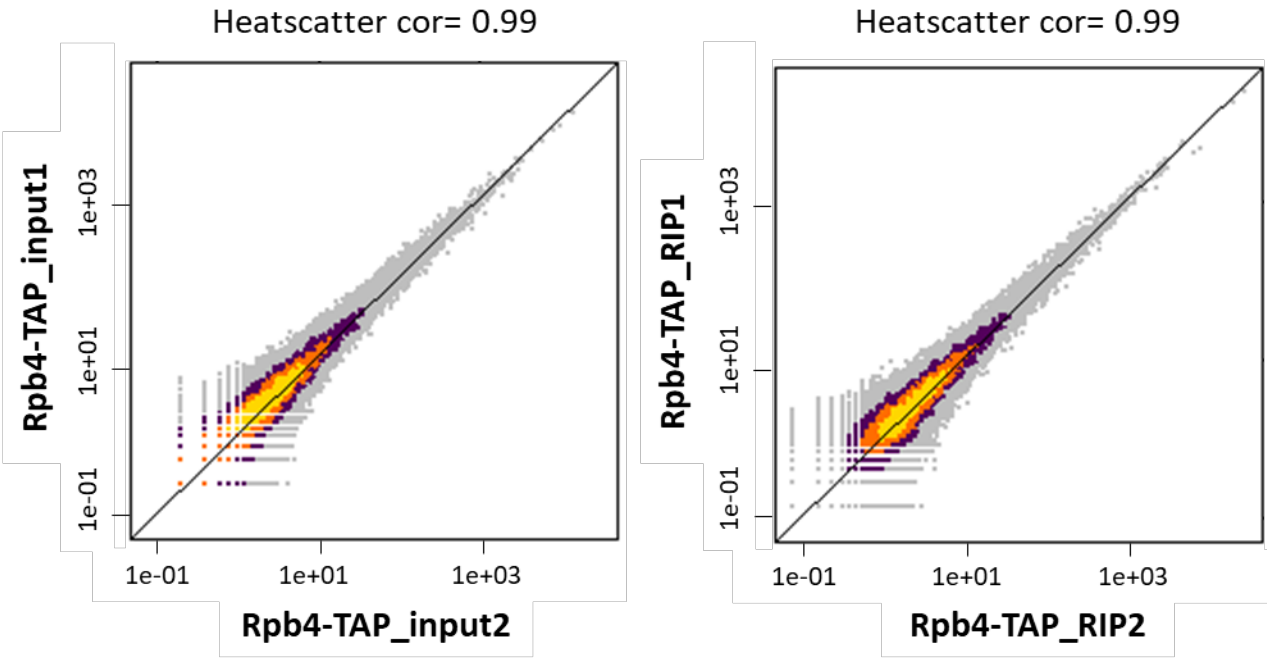


Figure S1

| | Rpb4-RIP cut-off: $Log_2FC>0$ (1458 mRNAs) | | Rpb4-RIP cut-off: $Log_2FC>0.58$ (778 mRNAs) | |
|---|---|---|---|---|
| | Common mRNAs | % of Rpb4-Puf3 common mRNAs over Rpb4-RIP data | Common mRNAs | % of Rpb4-Puf3 common mRNAs over Rpb4-RIP data |
| **RIP-Seq (Kershaw)** | 369 | 25.31 | 218 | 28.02 |
| **RIP-Seq (Gupta)** | 31 | 2.13 | 20 | 2.57 |
| **HITS-CLIP (Wilinski)** | 203 | 13.92 | 135 | 17.35 |
| **RIP-ChIP (Gerber)** | 139 | 9.53 | 107 | 13.75 |
| **PAR-CLIP (Freeber)** | 148 | 10.15 | 76 | 9.769 |
| **Puf3-bound mRNAs (in, at least, 3 datasets)** | 138 | 9.47 | 104 | 13.37 |

Figure S2

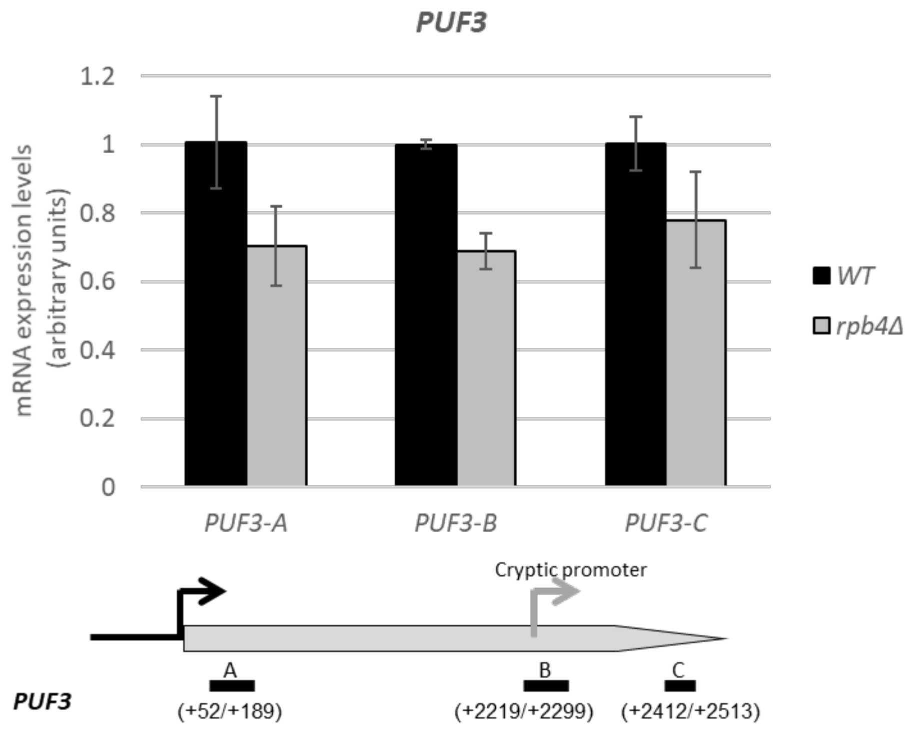
PUF3
1.2
1
0.8
0.6
0.4
0.2
0
mRNA expression levels (arbitrary units)
PUF3-A
PUF3-B
PUF3-C
WT
rpb4Δ
Cryptic promoter
A
B
C
PUF3
(+52/+189)
(+2219/+2299)
(+2412/+2513)

Figure S3

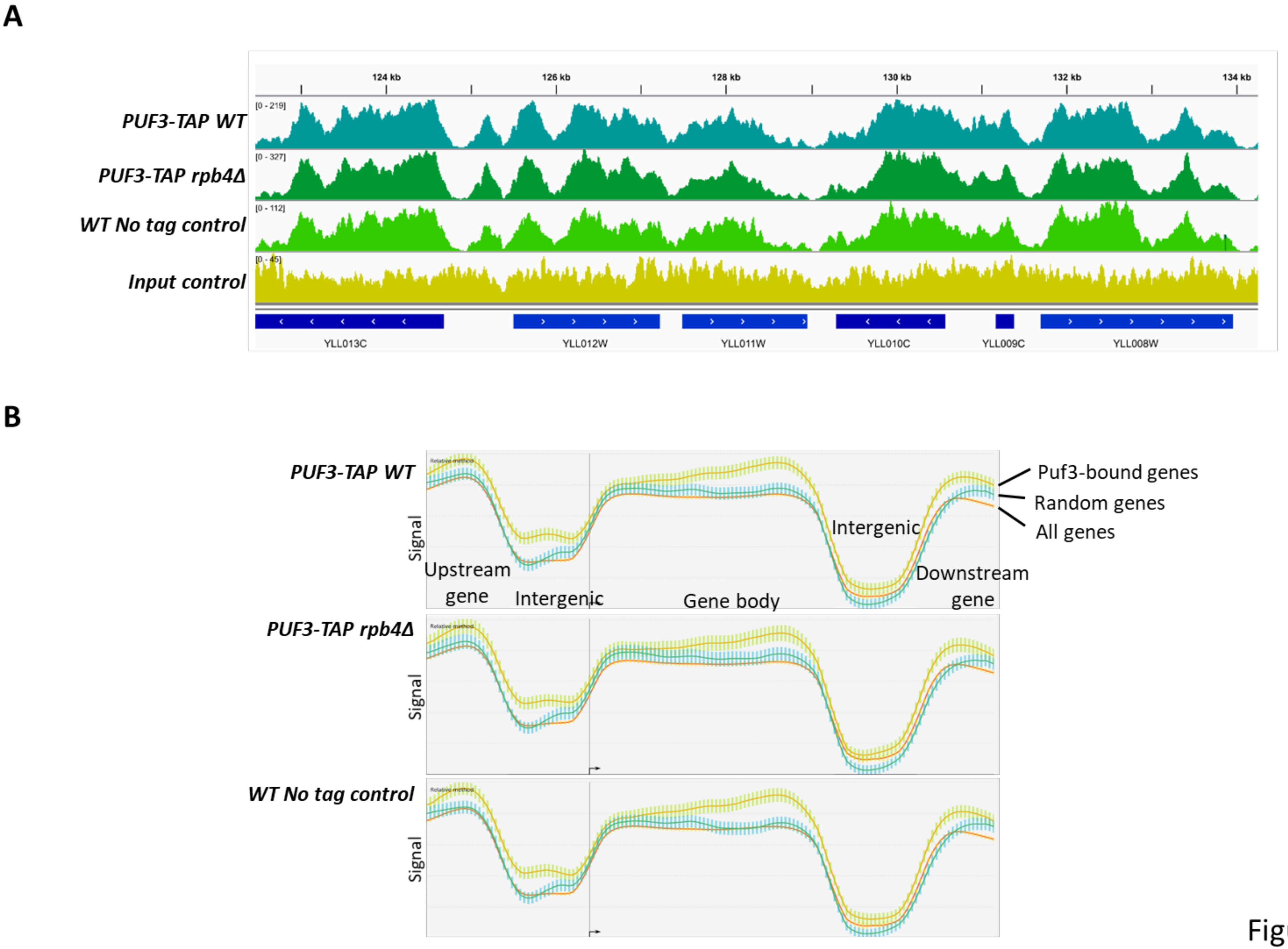
A
124 kb
126 kb
128 kb
130 kb
132 kb
134 kb
PUF3-TAP WT
[0 - 219]
PUF3-TAP rpb4Δ
[0 - 327]
WT No tag control
[0 - 112]
Input control
[0 - 45]
YLL013C
YLL012W
YLL011W
YLL010C
YLL009C
YLL008W
B
PUF3-TAP WT
PUF3-TAP rpb4Δ
WT No tag control
Signal
Upstream gene
Intergenic
Gene body
Intergenic
Downstream gene
Puf3-bound genes
Random genes
All genes

Figure S4

**Figure S1**: **Heatpairs showing a high correlation between two biological replicates**. Similarity between samples indicated by a numerical value, where 1 is the value of the greatest similarity, and scatterplots where the deviation of points from the trend line indicated variability between samples. Warmer colors indicate the higher density of the overlapping genes. Spearman rank correlation values are indicated (heatscatter cor).

**Figure S2**: **Comparison between the Rpb4-bound mRNAs from RIP-Seq (this study) and different datasets**. The Rpb4-bound mRNAs from the RIP-Seq identified in this study with different cut-offs ($log_2$ FC>0 and $Log_2$FC>0.58) were compared to the Rpb4-bound mRNAs from different datasets: RIP-Seq [7, 8], HITS-CLIP [9], RIP-CHIP [11] and PAR-CLIP [10].

**Figure S3: mRNA levels for the *PUF3* gene in the *rpb4Δ* mutant and the wild-type strains**. Cells were grown in YPD medium at 30ºC and analyzed by qRT-PCR. The *PUF3* mRNAs was measured with three pairs of different primers to detect differences in mRNA abundance that could be explained by the presence of an internal cryptic promoter. rRNA 18S was used as a normalizer. Data are shown as the average and standard deviation (SD) of at least three independent experiments.

**Figure S4: ChIP-Seq for Puf3**. **A)** Integrated genome browser (IGV) view of the mapped ChIP-Seq reads for the Puf3-TAP WT, Puf3-TAP *rpb4Δ* and WT no tag control samples, as well as input for the Puf3-TAP WT strain. Tracks are autoscaled as coverage differed among samples. *YLL009C* (*COX17*) is an example of a Puf3-regulated gene [12]. However, no difference was found in the peaks between any strains above this gene or others. **B)** Metagene profiles of the signal in Puf3-TAP WT, Puf3-TAP *rpb4Δ*, and WT no tag control ChIP over all the *S. cerevisiae* genes, Puf3-associated genes, or randomly subsampled genes plotted using the versatile aggregate profiler (VAP) [13].

1. Garrido-Godino AI, García-López MC, Navarro F. Correct assembly of RNA polymerase II Depends on the foot domain and Is required for multiple steps of transcription in *Saccharomyces cerevisiae*. Mol Cell Biol 2013; 33:3611-26.
2. Schulz D, Pirkl N, Lehmann E, Cramer P. Rpb4 subunit functions mainly in mRNA synthesis by RNA polymerase II. J Biol Chem 2014; 289:17446-52.
3. Duek L, Barkai O, Elran R, Adawi I, Choder M. Dissociation of Rpb4 from RNA polymerase II is important for yeast functionality. PLoS One 2018; 13:e0206161.
4. Laribee RN, Hosni-Ahmed A, Workman JJ, Chen H. Ccr4-Not regulates RNA polymerase I transcription and couples nutrient signaling to the control of ribosomal RNA biogenesis. PLoS Genet 2015; 11:e1005113.
5. Gavin AC, Bosche M, Krause R, Grandi P, Marzioch M, Bauer A, et al. Functional organization of the yeast proteome by systematic analysis of protein complexes. Nature 2002; 415:141-7.
6. Szklarczyk D, Morris JH, Cook H, Kuhn M, Wyder S, Simonovic M, et al. The STRING database in 2017: quality-controlled protein-protein association networks, made broadly accessible. Nucleic Acids Res 2017; 45:D362-D8.
7. Kershaw CJ, Costello JL, Talavera D, Rowe W, Castelli LM, Sims PF, et al. Integrated multi-omics analyses reveal the pleiotropic nature of the control of gene expression by Puf3p. Scientific reports 2015; 5:15518.
8. Gupta I, Clauder-Munster S, Klaus B, Jarvelin AI, Aiyar RS, Benes V, et al. Alternative polyadenylation diversifies post-transcriptional regulation by selective RNA-protein interactions. Mol Syst Biol 2014; 10:719.
9. Wilinski D, Buter N, Klocko AD, Lapointe CP, Selker EU, Gasch AP, et al. Recurrent rewiring and emergence of RNA regulatory networks. Proceedings of the National Academy of Sciences 2017; 114:E2816-E25.
10. Freeberg MA, Han T, Moresco JJ, Kong A, Yang YC, Lu ZJ, et al. Pervasive and dynamic protein binding sites of the mRNA transcriptome in *Saccharomyces cerevisiae*. Genome Biol 2013; 14:R13.
11. Gerber AP, Herschlag D, Brown PO. Extensive association of functionally and cytotopically related mRNAs with Puf family RNA-binding proteins in yeast. PLoS biology 2004; 2:e79.
12. Houshmandi SS, Olivas WM. Yeast Puf3 mutants reveal the complexity of Puf-RNA binding and identify a loop required for regulation of mRNA decay. RNA 2005; 11:1655-66.
13. Coulombe C, Poitras C, Nordell-Markovits A, Brunelle M, Lavoie MA, Robert F, et al. VAP: a versatile aggregate profiler for efficient genome-wide data representation and discovery. Nucleic Acids Res 2014; 42:W485-93.
